# Pure species discriminate against hybrids in the *Drosophila melanogaster* species subgroup

**DOI:** 10.1101/2020.07.22.214924

**Authors:** Antonio Serrato-Capuchina, Timothy D. Schwochert, Stephania Zhang, Baylee Roy, David Peede, Caleigh Koppelman, Daniel R. Matute

## Abstract

Introgression, the exchange of alleles between species, is a common event in nature. This transfer of alleles between species must happen through fertile hybrids. Characterizing the traits that cause defects in hybrids illuminate how and when gene flow is expected to occur. Inviability and sterility are extreme examples of fitness reductions but are not the only type of defects in hybrids. Some traits specific to hybrids are more subtle but are important to determine their fitness. In this report, we study whether F1 hybrids between two species pairs of *Drosophila* are as attractive as the parental species. We find that in both species pairs, the sexual attractiveness of the F1 hybrids is reduced and that pure species discriminate strongly against them. We also find that the cuticular hydrocarbon (CHC) profile of the hybrids is intermediate between the parental species. Perfuming experiments show that modifying the CHC profile of the hybrids to resemble pure species improves their chances of mating. Our results show that behavioral discrimination against hybrids might be an important component of the persistence of species that can hybridize.

## INTRODUCTION

Species are lineages that are genetically isolated from one another as a result of innate biological differences (Coyne and Orr 2004; Sobel et al. 2010; Harrison 2012; Nosil 2012; Harrison and Larson 2014). Identifying traits that encourage the initial partitioning of the genetic variation into clusters is critical for understanding how species form. In addition to understanding how new species arise, one of the main goals of speciation studies is to understand why species in secondary contact do not collapse into a single population. When speciation is recent, nascent species might have the chance to exchange genes and subsequently merge into a single genetic group. The traits that maintain potentially interbreeding lineages apart are key to understanding why some closely related species persist or collapse (Rosenblum et al. 2012).

Barriers to gene flow can be categorized on whether they occur before or after mating takes place and are deemed either prezygotic or postzygotic barriers, based on their occurrence relative to fertilization of the zygote (Dobzhansky 1937; Sobel et al. 2010). Prezygotic barriers include all the phenotypes of the pure species that preclude the formation of hybrids and range from habitat isolation to incompatibilities between gametes. Among the types of prezygotic isolation, behavioral isolation seems to be ubiquitous in animals (Janicke et al. 2019). Individuals recognize their species (conspecifics) and discriminate against individuals from other species (heterospecifics), based on the recognition of a combination of chemical, auditory, or visual cues (Cady et al. 2011; Vortman et al. 2013; Mérot et al. 2015). Postzygotic barriers include all fitness defects associated with hybrids (Orr and Presgraves 2000; Orr 2005). The most commonly studied forms of postzygotic isolation are hybrid inviability and sterility, in which hybrids fail to develop or even if they develop, do not produce viable gametes, respectively (Orr and Presgraves 2000; Orr 2005).

Other, less extreme, phenotypes can also cause postzygotic isolation. Insect hybrids often show a reduced ability to find proper substrates (Linn et al. 2004; Godoy-Herrera et al. 2005; Bendall et al. 2017; Turissini et al. 2017; Cooper et al. 2018). Hybrid birds show reduced ability to perform key tasks, exhibit a decrease in learning spatial tasks, and are worse than their parents at solving novel problems (Delmore and Irwin 2014; McQuillan et al. 2018). Hybrids in flowering plants are often less attractive to pollinators (Levin 1970; Campbell et al. 1997; Campbell 2004; Ippolito et al. 2004). Reductions of hybrid fitness exist along multiple axes more nuanced than complete infertility or inviability.

The sexual attractiveness of hybrids remains largely understudied (but see (Krebs 1990; Gottsberger and Mayer 2007, 2019; Svedin et al. 2008). Behavioral isolation, in the form of mate choice, can also be postzygotic and occur between species and the resulting hybrids (Gottsberger and Mayer 2007, 2019). Since fertile F1 hybrids constitute the bridge for genetic exchange between species, assessing the existence of any fitness defects in fertile hybrids constitutes a key component of understanding how much these individuals can facilitate gene exchange between species and determine whether introgression is favored in one of the directions of the cross. A natural prediction is that if species recognition depends on multiple traits, and hybrids show a combination of parental values in those traits, then hybrids might be less attractive to the parental genotypes, reducing hybrid fitness and the potential for introgression.

Cuticular hydrocarbons (CHCs) are fatty acid-derived apolar lipids that accumulate on the body cuticle of insects (reviewed in (Singer 1998; Ferveur 2005; Blomquist and Bagnères 2010)). The function of CHCs is twofold, affecting survival and reproduction (Chung and Carroll 2015). First, CHCs help regulate the osmotic balance within insect bodies, which makes them important for adaptation to water-limited areas. Second, CHCs are important for chemosensory communication among individuals. As a result, divergence in CHCs can lead to prezygotic isolation among species, both in terms of habitat separation and of mate choice (reviewed in (Smadja and Butlin 2009). Long-chained CHCs are usually more important in waterproofing, whereas shorter-chain CHCs, which tend to be more volatile CHCs, can be involved in sexual signaling over short distances (Hadley 1981; Gibbs 1998; Ferveur and Cobb 2010; Gibbs and Rajpurohit 2010). Longer chain CHCs can also act as contact pheromones in multiple insect species (Venard and Jallon 1980; Ingleby 2015). Closely-related species commonly differ in the composition of CHCs (Shirangi et al. 2009); these differences reduce the likelihood of matings between individuals from different species, effectively serving as a barrier to gene flow. Despite the robust research program reporting the differences in CHC composition between different species pairs (e.g., (Gleason et al. 2009; Sharma et al. 2012; Chung and Carroll 2015; Dembeck et al. 2015; Denis et al. 2015; Combs et al. 2018)), little is known regarding changes of CHC composition in F1 hybrids and how these changes might affect the attractiveness of hybrids to pure species individuals.

*Drosophila* species pairs in which hybridization yields fertile progeny, and show evidence of introgression in nature, are ideal systems to study barriers to gene flow that contribute to species persistence in nature. In this report, we studied whether interspecific hybrids are less attractive than pure-species counterparts to pure-species individuals. We focus on two species clades that produce fertile female progeny and show evidence of gene exchange in nature, the *Drosophila simulans* and *D. yakuba*-species complexes. The *Drosophila simulans*-species complex consists of three sister species: *D. simulans, D. sechellia*, and *D. mauritiana. Drosophila simulans* can produce fertile F1 females with both *D. sechellia* and *D. mauritiana*; the species triad diverged within the last 0.2 million years (Kliman et al. 2000; Schrider et al. 2018; Meany et al. 2019). All species pairs in this group show evidence of introgression (Garrigan et al. 2012; Brand et al. 2013; Meiklejohn et al. 2018; Schrider et al. 2018), and in the case of *D. simulans* and *D. sechellia*, the two species form a hybrid zone in the central islands of the Seychelles archipelago (Matute and Ayroles 2014). The *D. yakuba*-species complex is also composed of three species: *D. yakuba, D. santomea*, and *D. teissieri. Drosophila yakuba* and *D. santomea* diverged between 0.5 and 1 million years ago, while the dyad diverged from their sister *D. teissieri* approximately 3 million years ago (Bachtrog et al. 2006; Turissini and Matute 2017). Like the *D. simulans* complex, hybrid crosses involving *D. yakuba* with *D. santomea/D. teissieri* produce sterile males and fertile females (Lachaise et al. 2000; Coyne et al. 2004). Notably, *D. yakuba* forms stable hybrid zones with both *D. santomea* (Llopart 2005; Llopart et al. 2009; Matute 2010; Comeault et al. 2016) and *D. teissieri* (Cooper et al. 2018; Turissini and Matute 2017) in the Afronesian islands of São Tomé and Bioko, respectively.

In this study, we report that *Drosophila* hybrids—both male and female—are less likely than pure species individuals to be pursued and accepted in mating by pure species. The CHC composition of female hybrids is largely intermediate between parentals. Hybrid females perfumed as pure species show higher attractiveness to pure species females. Additionally, pure species perfumed as F1s show reduced attractiveness. These results suggest that hybrids are less sexually attractive than conspecifics, likely due to differences in CHC composition. Finally, we quantify CHC profiles of pure species and their hybrids to test how their pheromonal composition changes within the species complex. Our results suggest that nuanced fitness reductions in hybrids can be important to determine whether hybrids facilitate gene transfer between species in nature.

## METHODS

### Stocks

Our goal was to compare the attractiveness (mate choice tests) and CHC content of F1 hybrids to their pure species parents. To this end, we used isofemale lines for all our experiments. We used a single isofemale line for each of the four species we studied. Please note that there is variation between isofemale lines within species (Sharma et al. 2012; Denis et al. 2015) and extensive phenotypic plasticity in CHCs (Thomas and Simmons 2011; Rajpurohit et al. 2017; Otte et al. 2018). Details for each of these lines have been previously published. For *D. simulans*, we used the line Riaba, which was collected in 2009 on the island of Bioko (Serrato-Capuchina et al. 2020). For *D. mauritiana*, we used R50, a line collected on the island of Rodrigues in 2009 (Brand et al. 2013). For *D. yakuba*, we used ym5.02, a line collected in the midlands of the island of São Tomé in 2018. Finally, for *D. santomea*, we used Thena7, a line collected at the edge of Obó national park on the island of São Tomé (Comeault et al. 2016). All lines were kept in cornmeal 30mL vials.

### Fly rearing and virgin collection

During the experiments reported here, we kept all isofemale lines in 100ml plastic bottles with standard cornmeal/Karo/agar medium at room temperature. Once we saw larvae on the media, we transferred the adults to a different bottle and added a squirt of 0.5% V/V solution of propionic acid and a pupation substrate (Kimberly Clark, Kimwipes Delicate Task; Irving, TX) to the media. Approximately 10 days later, virgin pupae start eclosing, at which point we began collecting virgins. We cleared bottles every 8 hours and collected the flies that emerged during that period. This procedure ensured that flies had not mated, as they are not sexually mature. We separated flies by sex and kept them in sex-specific vials in groups of 20 individuals.

### Hybrid production

To make hybrids, we mixed a group of females and males (collected as described immediately above) in 30 mL vials with freshly yeasted food. All flies were 3-7 day old virgins. To increase the likelihood of mating, we mixed flies in a 1:2 female to male ratio. Vials were inspected every two days to see if there were larvae in them. Once we observed larvae in the vials, adults were transferred to a new vial, and the previous vial was tended with 0.5% propionic acid and a pupation substrate as described above. If, after a week, a vial had not produced progeny, the flies were transferred to a vial with fresh food. Virgin hybrids were collected in the same way described above and stored in sex-specific vials until further experimentation. To further ensure the identity of the hybrids, we extracted the testes of a subset of the F1 males for each cross (N=20) to score their fertility using methods previously published, namely, scoring for motile sperm (Turissini et al. 2015). F1 hybrids in all the possible crosses are sterile as they do not produce motile sperm (Coyne et al. 2004; Moehring et al. 2004; Turissini et al. 2015). In all instances reporting F1 hybrids, the genotype of the mother is listed first, and the genotype of the father, second.

### Mate choice tests

#### Effect of markings on female attractiveness

Our first set of experiments involved a setup with one male and four females for the male to choose. As a proxy of the attractiveness of each female in the vial, we measured the time the male spent courting each of the females. These male attractiveness experiments required labeling the females to distinguish them from each other. We marked the females in two different ways. First, we clipped their wings. To do this marking, we anesthetized flies at collection (∼ eight hours after hatching) and cut a nick in their wing in one of four ways: horizontal on the right wing, horizontal on the left wing, vertical on the right wing, or vertical on the right wing. Second, we placed the marked flies in dyed food two hours before matings to color their abdomens. We used three different colors and left one of the genotypes unlabeled for a total of four abdominal colors. We used a combination of both markings for a total of 16 potential combinations.

We studied the potential effect that these markings had on female attractiveness by doing mate choice experiments with a conspecific male and four females of the same genotype marked differently. To study the effect of single markings, we placed four females from the same genotype marked differently (either colored or clipped), and a conspecific male in a 30mL vial. We observed the group for one hour. We then let the male choose among the four females and scored the identity of the mated female. First, we assessed whether colored abdomens led to a change in attractiveness by labeling the females only with colored food (100 females per species). Second, we assessed whether the clipping procedure led to a change in attractiveness (100 females per species). For these two experiments (each with four types of marking), the proportion of chosen females should follow a 1:1:1:1 expectation as long as the marking treatment has no effect on the attractiveness of the females. We used Pearson’s *X*^2^ test to test these two hypotheses (function *chisq.test*, library ‘*stats’*, (R Core Team 2016)).

Finally, we studied whether the double-marking approach affected the female attractiveness in conspecific matings using mass matings. We labeled females with the two markings (color and clipping; 16 combinations). We then put 320 flies in a mesh cage (24.5 cm x 24.5 cm x 24.5 cm; www.bugdorm.com): 160 males and 10 females of each marking. We observed the cage for one hour and aspirated pairs that were mating and scored the marking of the male. We ran this experiment three times for each of the species for a total of 12 cages (480 males per species). Please note this experiment is different from our other choice experiments (described above) and since there are more males per cage than in a 30mL vial, there might be higher chance of reproductive interference among males (Matute 2014). Nonetheless, these results inform if any marking scheme grossly affects female attractiveness. We compared the proportions of mated females with the expectation of a uniform mating rate using a Pearson’s *X*^2^ test (function *chisq.test*, library ‘*stats’*, (R Core Team 2016).

Additionally, we ran a smaller experiment in which we studied the female attractiveness of doubly-marked females, measured as the time the males spent courting each female. We placed four marked females in a 30mL vial with a pure species conspecific male and scored the time that the male spent courting each type of female as described above. The marking of each female was randomly assigned. We did this for the four species and 24 replicates per species. The metric of attractiveness was scored by two different scorers. We only observed one trial at once. First, we assessed whether the scorer had an effect by fitting a linear model in which species and scorer were the effects and the observed time was the response (function *lm*, R library *stats*; (R Core Team 2016)). Since we found no strong effect of the observer, all further observations involved only one observer.

Second, and using the same dataset, we studied whether the double-markings had an effect on the attractiveness of each female. We fitted a linear model (function *lm*, R library *stats*; (R Core Team 2016)) where the proportion of time that each male spent courting each of the four types of females was the response and the two types of markings were the fixed effects. We included an interaction term between markings.

#### Female attractiveness in male choice experiments

To compare F1 hybrids vs pure-species female attractiveness, we used mate choice experiments in which a pure species virgin male had the choice of four different females, a virgin hybrid F1 female from each reciprocal cross, and one virgin female from each of the parental species in a 30mL vial with cornmeal food (i.e., five flies per mating assay). Since F1 females and pure species look similar, we marked them as described above. Even though our experiments show that these marking schemes have no effect on male choice (See below, Figures S1-S3), we randomized the genotype and the marking scheme to minimize any potential effect of the markings. The five flies (four females and one male) were placed in a vial within one minute and were not moved for the next two minutes. For the next 30 minutes, we observed what female the male approached and scored active courting behavior defined as the time that the male spent following, courting, and attempting to mount each type of female. We observed only one male at a time. We then scored an index of attractiveness for each female defined as:

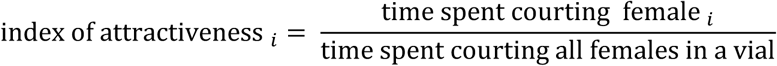

We observed 300 males from 15 different families per species for a total of 1,200 males (i.e., 20 males × 15 families × 4 species). To analyze the metric of attractiveness of each female (described above), and whether males courted conspecific and hybrid females at different rates, we compared the proportion of time that each male spent courting each of the four types of females using a linear mixed model (LMM; function *lme*, R library *nlme*; (Pinheiro J, Bates D, DebRoy S 2017)) where the identity of the female was a fixed effect and the block and marking were considered random effects. We also did posthoc pairwise comparisons using the function *lsmeans* (R library *lsmeans*, (Lenth and Hervé 2015; Lenth 2017)).

#### Male mating rates in non-choice experiments

Next, we studied the mean mating rates of hybrid males using a non-choice mating experiment (one female and one male). We used this design because it allowed us to assess whether a female accepted or rejected a given male while measuring the effort to mate excerpted by the male. To set up non-choice experiments, we placed a 4 day-old virgin female and a 4 day-old virgin male in a 30mL plastic vial with cornmeal food. We observed the pair for one hour. For each genotype of pure-species female, we did mating experiments with four types of males: conspecific, heterospecific, and the two reciprocal F1 males. We setup ten blocks of matings (i.e., days of experimentation) for each cross, each with 100 matings, for a total of 1,000 matings per cross. We watched 200 matings at a time, all of them with the same female genotype (50 of each type of cross). We did two matings in a single day which yielded 100 females for each type of cross per day. Since there were only two flies per vial (one female and one male), there was no need to mark either sex.

For each type of mating, we scored three characteristics of the mating. First, we recorded whether the female accepted the male. To compare the likelihood of mating, we fitted a logistic regression. Females that mated were considered successes, females that did not were considered failures. The only fixed effect was the genotype of the male while the experimental block was considered a random effect. We used the function glmer (library *lme4* (Bates et al. 2015)) for these analyses. To assess significant of the fixed effect, we used a type-III ANOVA (function *Anova*, library *car*; (Fox and Sanford 2011; Fox and Weisberg 2019)). We also measured the significance of the male effect using a likelihood ratio test comparing the model described above with one of no male effect (function *lrtest*, library *lmtest;* (Hothorn et al. 2011)). We compared the proportion of accepted males in the different crosses using a Tukey test using the function *lsmeans* (library *lmtest*, (Hothorn et al. 2011)).

Second, we recorded the copulation latency for the different types of crosses, which also serves as an additional proxy of male attractiveness (i.e., more attractive males have shorter latencies, (Ejima and Griffith 2007)). Since the likelihood of the different matings differs, the number of observations differs among cross types (See Results). To compare latency and duration among cross-type, we used the function *lm* (library *stats*, (R Core Team 2016)) and fitted a one-way ANOVA where the latency was the response, and the cross was the only fixed factor. We also did posthoc pairwise comparisons using a Tukey HSD test (function *glht, library ‘multcomp’* (Hothorn et al. 2008)). We used an identical approach to study heterogeneity in copulation duration.

We also recorded whether the male was courting the female every two minutes (the time that it took to inspect the 200 males in the assay). Every time window in which the male was observed, courting was categorized as an effort to mate. To study male effort by genotype, we focused on pairs that mated. Since one might observe higher efforts on longer matings (i.e., higher latencies), we used the number of efforts per unit of time, a ratio between the number of efforts and the mating latency. We compared this metric across genotypes with a linear model where the metric was the response, and the type of cross was the fixed factor (function *lm*, library *stats*).

### CHC quantification

#### Studied CHCs

For *D. simulans, D. mauritiana* and their hybrids, we measured the concentrations of n-Heneicosane, 11-*cis*-Vaccenyl Acetate, Tricosane, 7(Z)-Tricosene, 7-Pentacosene, 7(Z),11(Z)-Heptacosadiene, and 7(Z)-Nonacosadiene. For *D. yakuba, D. santomea*, and their hybrids, we measured the same CHCs. These CHCs encompass the primary CHC composition in both the *simulans* (Sharma et al. 2012; Ingleby et al. 2013) and *yakuba* species complex (Mas and Jallon 2005; Denis et al. 2015).

#### CHC standard curves

To quantify the seven CHCs listed above, we purchased standards of the seven compounds. The catalog numbers are listed in Table S1. We did gas chromatography (GC) using an Agilent 7820A gas chromatography system equipped with a FID detected and a J&W Scientific cyclosil-B column (30 m × 0.25 mm ID × 0.25 µm film) to characterize the elution time of the standards. GC provides 1) the retention time of each CHC and 2) the peak integration ratio between the known quantities of the target CHC and that of internal standard permitting quantification of target CHCs from fly extracts. First, we measured the retention times for each of the CHCs (Table S1). This allowed us to identify specific CHC compounds in the fly extracts based on their retention time. Second, we diluted each of the compounds to concentrations of 150µM, 100 µM, 75 µM, 50 µM, and 25 µM using hexacosane (1mM) as an internal standard. We measured the signal ratio between the target CHC and that of the internal standard. For each compound, we fit a linear model using the function *lm* (library ‘*stats’*) with the concentration of the CHC as the response and the ratio of the peak height to the internal standard as the sole continuous factor. Figure S4 shows the seven regressions for each CHC.

#### CHC extraction from individual flies

CHCs from single virgin flies were extracted by placing individuals (aged 4-7 days) in a glass disposable culture tube and submerging in 1 mL of a solution of heptane and hexacosane (internal standard; mM) for 3 minutes with light shaking. The extract was filtered through glass wool prior to GC analysis. All extractions were completed between 2:00 pm and 4:00 pm, with GC analysis taking place as quickly as possible following the extraction procedure. Measurements were done in the same GC machine described above. The method used to separate CHCs present in fly extracts consisted of holding the GC oven at 150 °C for 5 min, then ramped at 5°C/min, held for 10 min, then ramped again at 10 °C/min, and held for 15 min. The number of samples for each genotype ranged between six and twenty-one and is listed in Table S2.

#### CHC quantification

We integrated the peaks using the Agilent 7280A software for each GC graph from individual extractions, and transformed the area-under-curve (AUC) across each corresponding retention time to CHC amount using the slopes of the calibration regressions, described above in ‘<u>CHC Standard curves</u>’. The conversion followed the transformation:

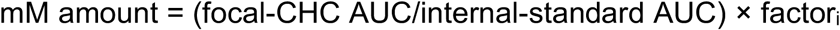

where factor_i_ represents the slope for each given CHC.

#### Analyses

To compare the CHC blends among pure species and F1 hybrids in each species group, we generated a principal component analysis (PCA) for each species group. We used the function *prcomp* (library *stats*) to calculate the PCA loadings and visualized the results with the function *fviz_pca_ind* (library *factoextra*, (Kassambara and Mundt 2017)). The distribution of each genotype was plotted using the option *ellipse.type* with a multinomial distribution. In both cases, PC1 explained the vast majority of the variance (see results), and we only used that PC to study heterogeneity among genotypes. We fitted a One-way ANOVA with PC1 as a response and genotype as the only factor using the function *lm* (library ‘*stats’*, (R Core Team 2016)). We then performed pairwise comparisons using a Tukey HSD test using the function *glht* (library *‘multcomp’*, (Hothorn et al. 2008)).

### Perfuming assays

Hybrids show lower success mating compared to pure species individuals, and show CHC profiles that differ from both parentals (See results). We tested whether there was a connection between these two results by changing the CHC profile, through perfuming, of hybrid and pure-species flies and then measuring their attractiveness. Perfuming consisted of placing a single focal female in a vial with 10 perfuming-females for 4 days, allowing time for the CHC profiles to homogenize. We describe these two sets of experiments as follows.

First, we did choice experiments with perfumed flies involving hybrid females. We focused on hybrid females as, unlike hybrid males, they are fertile and thus can serve as a bridge for gene exchange between species. The choice experiments involved a single pure-species male, and he had the choice to mate with one of three hybrid females (i.e., all with the same genotype), one that was not perfumed, one that was perfumed with one of the parents, and one with the other parent. We only used one of the reciprocal crosses per species pair (either ♀*yak/*♂*san* or ♀*sim/*♂*mau*) as they are much easier to produce than the reciprocal direction (Yukilevich 2012; Lachaise et al. 1986; Turissini et al. 2018). All perfuming experiments are similar, so we only describe one of them. To assess whether perfuming F1 (♀ *sim* × ♂*mau*) females changed their attractiveness to *D. mauritiana* males, we perfumed F1 (♀*sim*/♂*mau*) females with *D. mauritiana* females, F1 (♀*sim*/♂*mau*) females with *D. simulans* females, and F1 (♀*sim*/♂*mau*) females with other F1 (♀*sim*/♂*mau*) females. To perfume a F1 (♀*sim*/ ♂*mau*) female with *D. mauritiana* CHCs, we placed a single five day-old F1 (♀*sim*/♂ *mau*) female with 10 *D. mauritiana* females from the same sex for four days. The F1 (♀*sim*/♂*mau*) female perfumed with *D. simulans* was placed with *D. simulans* females instead but followed the same approach. The third female, was a F1 (♀*sim*/♂*mau*) that was ‘perfumed’ with other F1 CHCs by raising it with other ten F1 females of the same genotype. This procedure should not change the CHC profile of the female, but it exposes the focal female to the same rearing density of the other perfumed females. Females were marked by feeding them colored food and clipping their wings (see above). The three types of females were marked by labeling their abdomens and clipping their wings as described above. After four days, we removed each of the focal females from their ‘perfuming vials’ by aspiration (no CO_2_ anesthesia) and placed them in an 30mL vial with fly food. We also added a virgin pure species male with the three perfumed F1 females. We watched the vial for one hour and scored the identity of the female the male chose for mating. The expectation was that if perfuming had no effect on the attractiveness of the hybrid females, then the males should choose randomly and the choice should follow a 1:1:1 ratio. On the other hand, if the CHC blend on the hybrids reduces their attractiveness, then perfuming them like pure species should lead to an increase in their attractiveness (i.e., they should be more likely to be chosen by pure species male). We observed 50 flies per genotype in each block (i.e., experiments ran on the same day) and did six blocks per type of assay for a total of 300 per male genotype.

Second, we did similar experiments for each of the pure species and studied whether perfuming pure-species females with heterospecific, or hybrid CHC blends, reduced their attractiveness. We placed a pure species male with three conspecific females, one that was perfumed with her conspecifics, one that was perfumed with F1 hybrids, and one that was perfumed with the other species. The approaches of this set of experiments are identical to the ones described above for the F1 hybrids. The expectation was that if the heterospecifics or hybrid CHC blends are less attractive than the conspecific blend, then the females perfumed with these blends should be less attractive. We did 50 replicates for each type of male per block and six blocks, which lead to 300 observations per male genotype.

The analyses for both perfuming experiments, hybrids, and pure-species, are the same. To evaluate the 1:1:1 expectation, we used a Pearson’s *X*^2^ test (function *chisq.test*, library ‘*stats’*, (R Core Team 2016)). If the perfuming affected the outcome of the mating (i.e., the mated female), then the ratio of mated females from each treatment will differ from 1:1:1. To evaluate which pairs differed from each other, we used an Approximative Two-Sample Fisher-Pitman Permutation Test (function ‘*oneway_test*’, library ‘*coin’*; (Hothorn et al. 2006)).

A sample of these perfumed flies—from both perfumed pure species and perfumed F1s—was scored for CHC profiles as described above (See <u>CHC</u> <u>quantification</u>). The number of samples for each treatment ranged between four and twenty-one and is listed in Table S2.

## RESULTS

### Pure species discriminate against hybrids

First, we studied whether marking females had an effect on mate choice. We found that individual markings have no effect on the male choice (Figures S1-S2, Tables S3-S4). Double markings caused no deviations from the expectation of uniform male choice in mass matings either (in all cases *X*^*2*^ < 6.096, df = 15, P > 0.9, Table S5).

Since markings did not affect female attractiveness, we used them in matings where males had the choice of conspecific, heterospecific, and reciprocal hybrid females. Our goal was to determine whether pure species males discriminated against hybrid females. For all the four genotypes, the proportion of assays that yielded a mated male was over 80% (Figure 1A-D). As expected, pure species males from any of the four assayed species overwhelmingly preferred females from their own species over any other type of female—including hybrids—. In all assays, over 95% of the mated males chose conspecific females. The preference for conspecifics is consistent with previous results, which suggest that males show a strong preference for conspecific females and discriminate against heterospecific females (Shahandeh et al. 2018).

**FIGURE 1.**
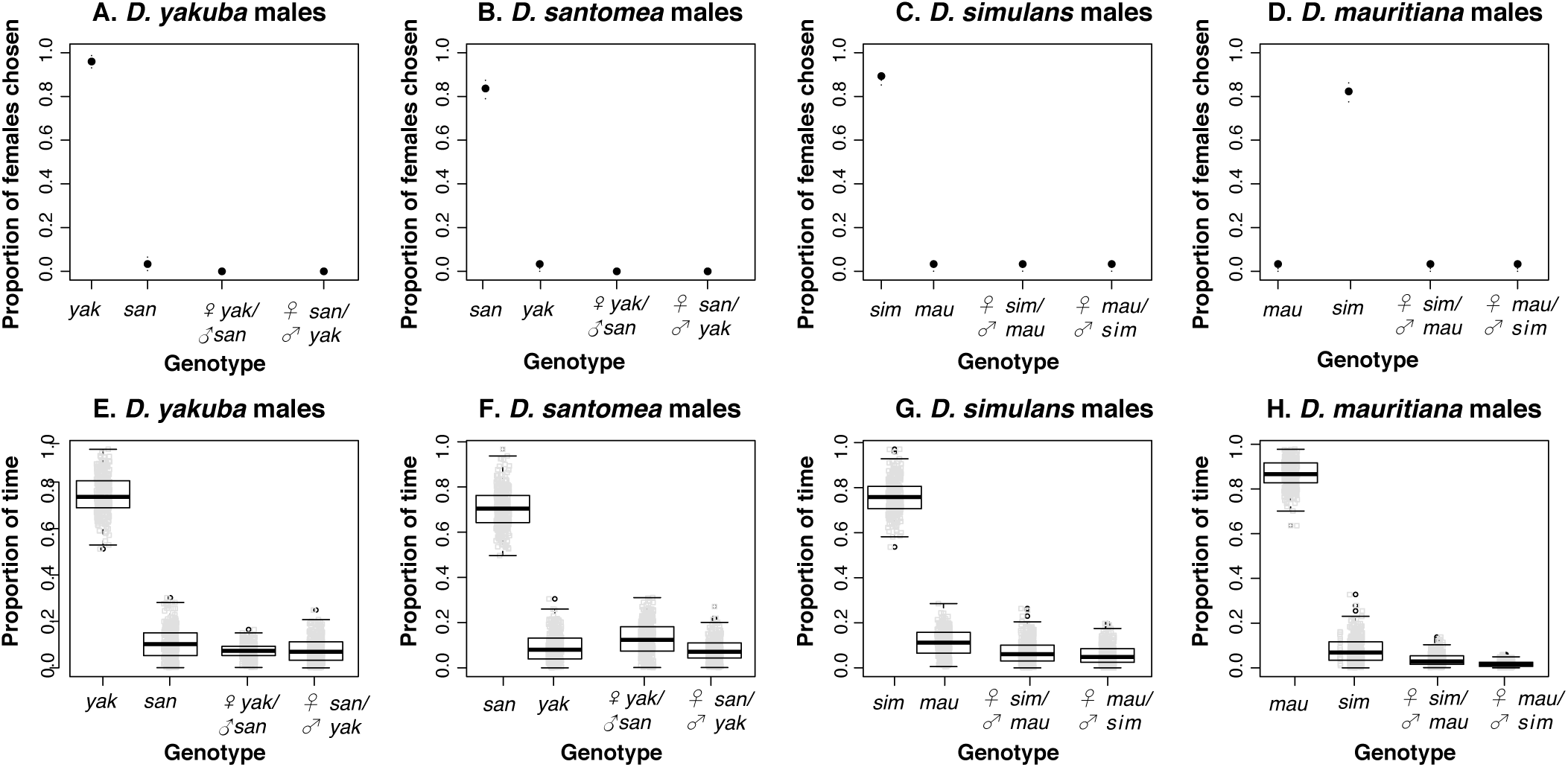
Pure species males discriminate against heterospecific and hybrid females in mate choice experiments. **A-D.** Proportion of males that chose a pure-species conspecific female in mating experiments where they had the choice of mating with conspecifics, heterospecfics, or F1 hybrid females. The black dot represents the proportion of conspecific matings (N=300) and the bars show the 95% confidence intervals calculated as Bayesian binomial intervals. **A.** *D. yakuba* males **B.** *D. santomea* males **C.** *D. simulans* males **D.** *D. mauritiana* males. **E-H.** Female attractiveness measurements in choice mating experiments. The proxy of female attractiveness is the proportion of time that a male spends courting each type of female relative to the total time that male spent courting all females. **E.** *D. yakuba* males **F.** *D. santomea yakuba* males **G.** *D. simulans* males **H.** *D. mauritiana* males.

Besides the outcome of the matings in mass matings, we also scored the effort males spent courting each type of female when they have four females to choose from. In choice experiments where the four females were conspecifics, we found no effect of the markings (Table S6, Figure S3) or the scorer (Figure S5) on female attractiveness. In experiments where males had the choice of mating with conspecific, heterospecific, and hybrid females, we found that the amount of time pure-species males spent courting each type of female differed depending on the type of female (Figure 1;LMM, female genotype effect F_3,957_ = 6,021.967, P <1 × 10^−10^ for all four types of males). Males spend much more time courting conspecific females than any other genotype (Figure 1E-H, Table S7). In general, F1 hybrid females were slightly less attractive than heterospecific females. Differences between reciprocal F1s were minimal (Figure 1, Table S7). These joint results indicate that pure species parental males are less likely to mate with F1 females if they have the choice of mating with a conspecific or heterospecific and that discrimination against hybrid females might act as an important component of reproductive isolation.

Next, we studied the frequency of mating of pure species females with conspecific, heterospecific, and hybrid males in non-choice mate experiments. For all female genotypes, the frequency of matings with heterospecific or hybrid males is much lower than the frequency of matings with conspecific males (Figure 2; LMM male genotype effect: LRT > 1,747.8; P < 1 × 10^−10^ in all cases, Table S8). In *D. yakuba, D. santomea*, and *D. simulans*, matings with hybrid males are less likely to occur than matings with heterospecific males (in *D. mauritiana* they are equally likely; Table S9). Lower rates of mating between pure-species females and hybrid males can be interpreted as lower male attractiveness, lower interest in matings by the males, or a combination of both. We measured a proxy of the effort invested by conspecific, heterospecific, and hybrid males in each type of cross in cases where mating took place. We find that the effort from males in heterospecific matings is generally lower than that in conspecific crosses (Figure 3). The effort in mating from hybrid males is similar to that shown by heterospecific males (Figure 3, Table S10). These results suggest that hybrid males have a lower interest in mating with either type of pure species female than pure species males do. They also show that even though hybrid males try to mate at similar rates as males in heterospecific crosses, their success rates are lower; thus, suggesting that the attractiveness of hybrid males is lower than that of either pure species males.

**FIGURE 2.**
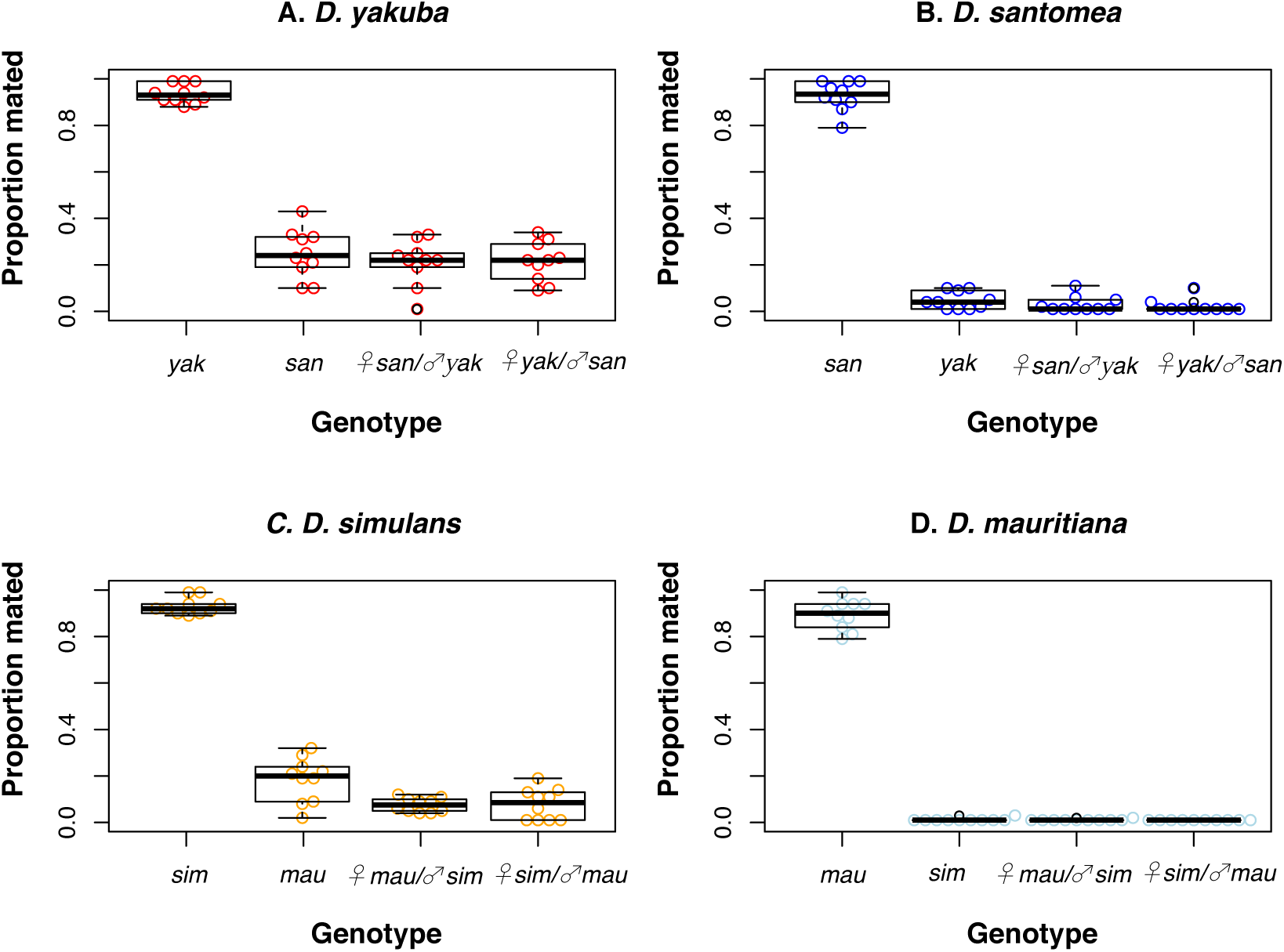
Pure species females engage in matings with heterospecific and hybrid males more rarely than they do with conspecific males in no-choice mating experiments. Proportion mated (y-axis) indicates the proportion of matings that led to a copulation (N=1,000). **A.** No-choice experiments with a *D. yakuba* female and one of four types of males from different genotypes (*D. yakuba, D. santomea*, F1 ♀*san/***♂***yak* hybrid, F1 ♀*yak/***♂***san* hybrid). **B.** No-choice experiments with a *D. santomea* female and one of four types of males from different genotypes (*D. santomea, D. yakuba*, F1 ♀*san/***♂***yak* hybrid, F1 ♀*yak/***♂***san* hybrid). **C.** No-choice experiments with a *D. simulans* female and one of four types of males from different genotypes (*D. simulans, D. mauritiana*, F1 ♀*mau/***♂***sim* hybrid, F1 ♀*sim/***♂***mau* hybrid). **D.** No-choice experiments with a *D. mauritiana* female and one of four types of males from different genotypes (*D. mauritiana, D. simulans*, F1 ♀*mau/***♂***sim* hybrid, F1 ♀*sim/***♂***mau* hybrid). Pairwise comparisons are shown in Table S9.

**FIGURE 3.**
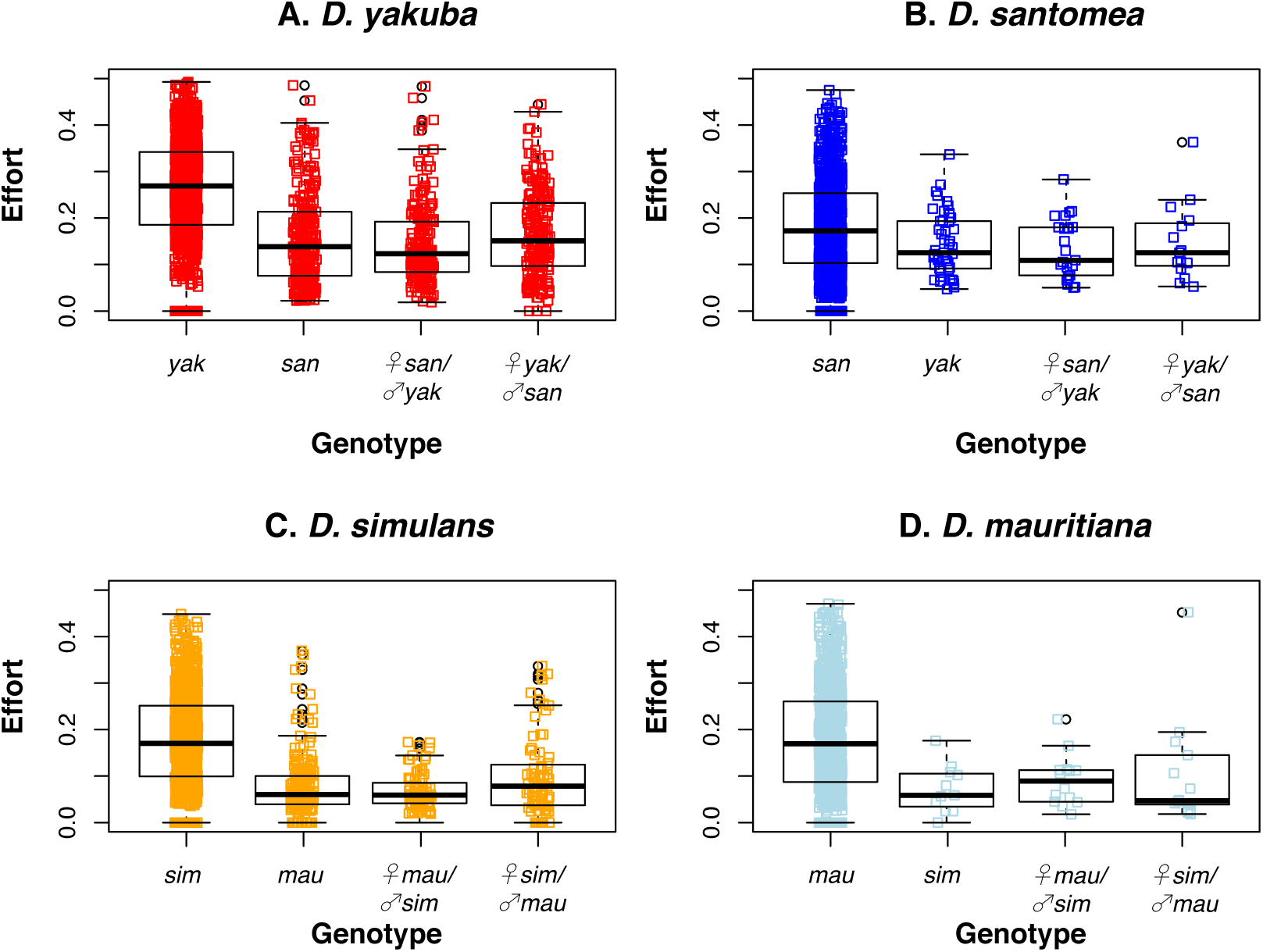
Males put more effort in matings with conspecific matings than in matings with heterospecific and hybrid females in non-choice experiments. The proxy of effort is the number of time windows (scored every 2 mins) in which the males were courting the female divided by the mating latency. **A.** No-choice experiments with a *D. yakuba* female and one of four types of males from different genotypes (*D. yakuba, D. santomea*, F1 ♀*san/♂yak* hybrid, F1 ♀*yak/♂san* hybrid). **B.** No-choice experiments with a *D. santomea* female and one of four types of males from different genotypes (*D. santomea, D. yakuba*, F1 ♀*san/♂yak* hybrid, F1 ♀*yak/♂san* hybrid). **C.** No-choice experiments with a *D. simulans* female and one of four types of males from different genotypes (*D. simulans, D. mauritiana*, F1 ♀*mau/♂sim* hybrid, F1 ♀*sim/♂mau* hybrid). **D.** No-choice experiments with a *D. mauritiana* female and one of four types of males from different genotypes (*D. mauritiana, D. simulans*, F1 ♀*mau/♂sim* hybrid, F1 ♀*sim/♂mau* hybrid). Pairwise comparisons are shown in Table S10.

We measured two additional characteristics of mating in these no choice experiments: copulation latency and copulation duration. When females mate with heterospecific or hybrid males in non-choice experiments, the matings take much longer to start than in non-choice conspecific matings. Latency is similar in matings with hybrids or with heterospecifics (Table 1). Mating duration is also longer in conspecific than in heterospecific of hybrid-male matings (Table S11). Altogether, these results are in line with the idea that the reduced mating rates of hybrid males are the result of lower interest in mating from the hybrids (behavioral sterility) and female discrimination against them. Since hybrid males in the two studied pairs are sterile, they cannot interbreed with the parental females; the fact that they are less likely to be accepted by the females is much less relevant as a reproductive barrier than their complete hybrid sterility. Nonetheless, this result is qualitatively similar to the pattern for female acceptance rates to different genotypes of males and suggests that the discrimination of pure species against hybrids is a phenomenon that applies to both sexes.

**TABLE 1.** Copulation latency in matings with conspecific, heterospecific, and hybrid males in no-choice experiments. Matings with heterospecific and hybrid males take longer to occur than conspecific matings. *N* represents the number of mated pairs (out of 1,000 attempts) used for the analyses. All the means (percentage of females mated) and standard deviations (SD). The last four columns show pairwise comparisons as 4 × 4 matrices for each cross. The lower triangular matrix shows the t value from a multiple comparisons of means using Tukey contrasts. The upper triangular matrix shows the P-value associated to the comparison. All p-values were adjusted for multiple comparisons.

| ***D. yakuba* females** |  |  |  |  |  |  |
| --- | --- | --- | --- | --- | --- | --- |
| Male genotype | N | Mean (SD)/min | Pairwise comparisons |  |  |  |
|  |  |  | *D. yakuba* | *D. santomea* | F1 (♀ *yak* × ♂ *san*) | F1 (♀ *san* × ♂ *yak*) |
| *D. yakuba* | 939 | 11.300 (4.594) | * | < 0.001 | <0.001 | <0.001 |
| *D. santomea* | 247 | 31.293 (10.865) | 33.361 | * | <0.001 | 0.912 |
| F1 (♀ *yak* × ♂ *san*) | 214 | 27.864 (12.820) | 26.091 | 4.382 | * | 0.002 |
| F1 (♀ *san* × ♂ *yak*) | 210 | 30.780 (11.771) | 30.450 | 0.652 | 3.583 | * |
| ***D. santomea* females** |  |  |  |  |  |  |
| Male genotype | N | Mean (SD) /min | Pairwise comparisons |  |  |  |
|  |  |  | *D. santomea* | *D. yakuba* | F1 (♀ *yak* × ♂ *san*) | F1 (♀ *san* × ♂ *yak*) |
| *D. santomea* | 928 | 24.480 (10.450) | * | < 0.001 | 0.006 | < 0.001 |

| *D. yakuba* | 45 | 40.694<br>(7.993) | 10.311 | * | 0.062 | 0.557 |
| --- | --- | --- | --- | --- | --- | --- |
| F1 (♀ *yak* × ♂ *san*) | 15 | 33.161<br>(10.311) | 3.238 | 2.452 | * | 0.559 |
| F1 (♀ *san* × ♂ *yak*) | 25 | 37.429<br>(8.037) | 6.202 | 1.271 | 1.268 | * |
| ***D. simulans* females** |  |  |  |  |  |  |
| Male genotype | N | Mean (SD) /min | Pairwise comparisons |  |  |  |
|  |  |  | *D. simulans* | *D. mauritiana* | F1 (♀ *sim* × ♂ *mau*) | F1 (♀ *mau* × ♂ *sim*) |
| *D. simulans* | 931 | 14.173<br>(5.829) | * | <0.001 | <0.001 | <0.001 |
| *D. mauritiana* | 185 | 30.857<br>(12.961) | 32.634 | * | 0.055 | <0.001 |
| F1 (♀ *sim* × ♂ *mau*) | 75 | 33.358<br>(10.168) | 19.032 | 2.502 | * | <0.001 |
| F1 (♀ *mau* × ♂ *sim*) | 75 | 40.460<br>(7.587) | 29.986 | 7.102 | 8.052 | * |
| ***D. mauritiana* females** |  |  |  |  |  |  |
| Male genotype | N | Mean (SD) /min | Pairwise comparisons |  |  |  |
|  |  |  | *D. mauritiana* | *D. simulans* | F1 (♀ *sim* × ♂ *mau*) | F1 (♀ *mau* × ♂ *sim*) |
| *D. mauritiana* | 894 | 21.038<br>(10.117) | * | <0.001 | <0.001 | <0.001 |
| *D. simulans* | 11 | 40.087<br>(6.053) | 6.179 | * | 0.997 | 0.952 |
| F1 (♀ *sim* × ♂ *mau*) | 14 | 39.249<br>(14.291) | 6.653 | 0.205 | * | 0.860 |
| F1 (♀ *mau* × ♂ *sim*) | 14 | 42.182<br>(10.798) | 7.725 | 0.512 | 0.764 | * |

### Hybrids show different CHC profiles than pure species

We measured the CHC composition of pure species and hybrids for two of the species pairs we studied, *D. yakuba/D. santomea* and *D. simulans/D. mauritiana*. We focused on females as they are fertile and can produce advanced intercrosses, whereas males cannot. Since the effect of CHCs in mating attractiveness is a joint one across multiple CHCs (Mas and Jallon 2005; Liimatainen and Jallon 2007; Grillet et al. 2012) and not one of individual CHCs, we plotted the distribution of the parental species and each of the F1 hybrids in a principal component analysis for each species pair (Figure 4). Tables S12 and S13 show the PCA loadings, and Figures S6 and S7 show the eigenvectors. The results for both species pairs are similar. In the case of *D. yakuba/D. santomea* females, we found that PC1 explains 97% of the variance. PC2 explains 1.4% of the variance. In the case of *D. simulans/D. mauritiana* females, we found that PC1 explains 76% of the variance. PC2 explains 22.5% of the variation. For both species pairs, PC1 differentiates between the two pure species; while PC2 seems to be associated with variance within genotypes. The CHC profile of the pure species is mostly disjointed in both species pairs. F1 hybrid females appear mostly as an intermediate between the two parental species, although some individuals seem to show transgressive patterns of segregation, consistent with other observations in other *Drosophila* hybrids (Coyne et al. 1994; Gleason and Ritchie 2004).

**FIGURE 4.**
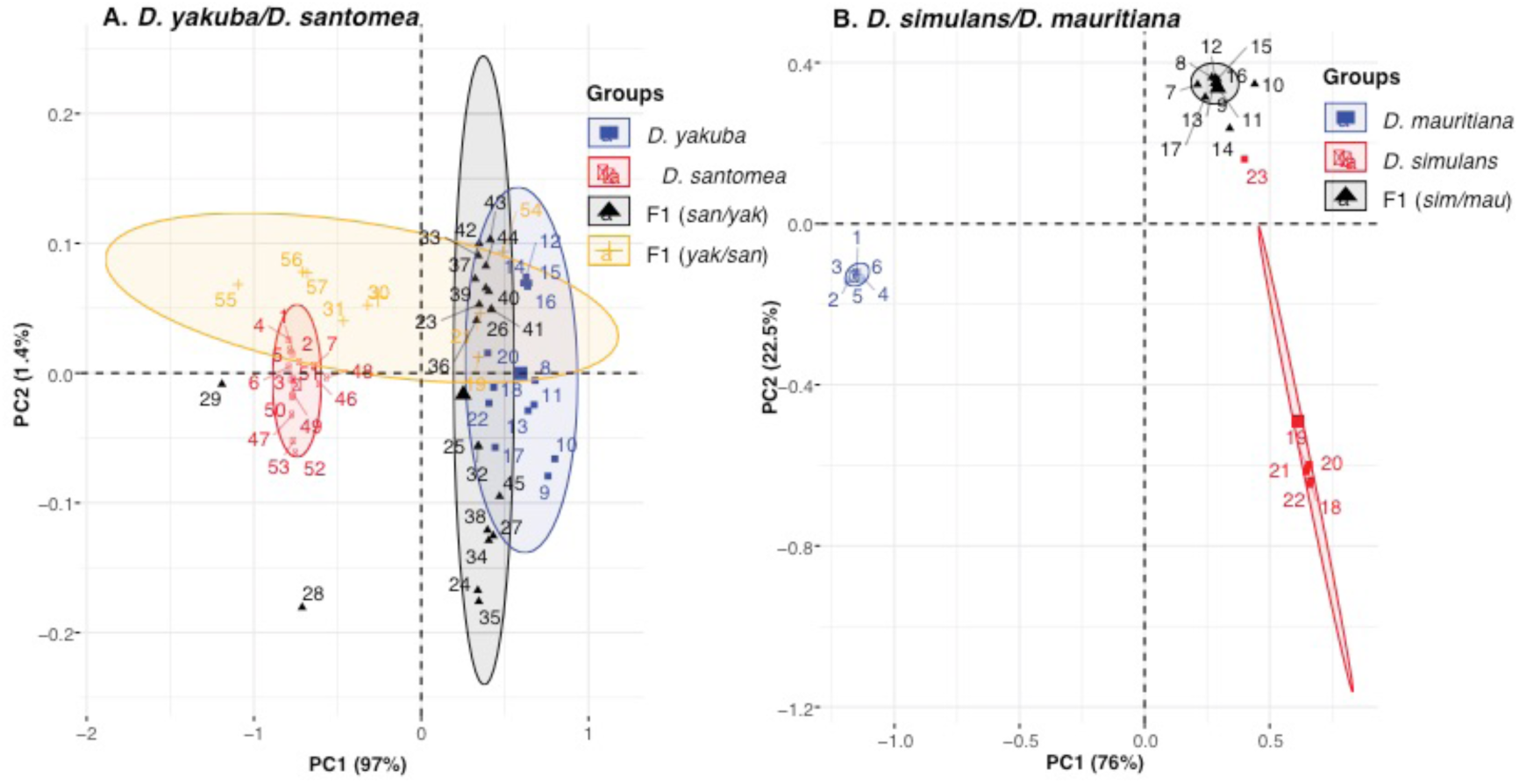
Principal component analysis (PCA) of cuticular hydrocarbon (CHC) profiles of pure species and hybrid females. Ellipses indicate a multinomial distribution of the data, variance explained by each PC is given in parentheses. PCA is based on the quantity of four CHCs for *D. yakuba* and *D. santomea* and six for *D. simulans* and *D. mauritiana* (see methods). **A.** *D. yakuba, D. santomea*, and reciprocal F1 hybrids **B.** *D. simulans, D. mauritiana*, and F1 ♀*sim/♂mau* hybrids.

Next, we studied whether there is variation in PC1 among genotypes. We found extensive heterogeneity among genotypes in both species pairs (Table 2). *Drosophila yakuba* and *D. santomea* differed, suggesting differences in their CHC blend. This is consistent with previous reports which showed differentiation in the CHC blends between this two species (Mas and Jallon 2005). The two reciprocal F1 females, F1 (♀*yak*/♂*san*) and F1 (♀*san*/♂*yak*) differ from all other genotypes (Table 2). F1 (♀*yak*/ ♂*san*) are broadly distributed along PC1 and PC2. Results were similar when we studied PC1 in the *sim/mau* genotypes (Table 2). (Note that we did not measure CHCs in F1 (♀*mau*/♂*sim*) hybrids.) All measured genotypes differ from each other (Table 2). F1 (♀*sim*/♂*mau*) hybrids showed a large spread on PC1. These results and the distribution of the CHC blend in the PCAs indicate that there are some differences between hybrids and pure species, but also that—at least some— F1s show a large variance in their CHC blends.

**TABLE 2.** Pure species and F1 hybrids tend to show differences in their joint CHC profile in the two studies species pairs. We performed pairwise comparisons using a Tukey test following a One-way ANOVA. The lower triangular matrix shows the t value from a multiple comparisons of means using Tukey contrasts. The upper triangular matrix shows the P-value associated to the comparison. Please note that we limited our analyses to PC1, because that PC explains over 75% of the variance in both species pairs. Figure 4 shows a representation of the same results.

| *D. yakuba*/ *D. santomea*: $F_{3,53} = 42.185$, $P < 1 \times 10^{-10}$ | | | | |
| --- | --- | --- | --- | --- |
| Pairwise comparisons |  |  |  |  |
| Genotype 1\Genotype 2 | *D. yakuba* | *D. santomea* | F1 (♀ *yak* × ♂ *san*) | F1 (♀ *san* × ♂ *yak*) |
| *D. yakuba* | * | < 0.001 | < 0.001 | 0.0269 |
| *D. santomea* | 10.345 | * | 0.0129 | < 0.001 |
| F1 (♀ *yak* × ♂ *san*) | 5.645 | 3.160 | * | 0.003 |
| F1 (♀ *san* × ♂ *yak*) | 2.892 | 8.576 | 3.648 | * |
| *D. simulans*/ *D. mauritiana*: $F_{2,20} = 1,225.1$, $P < 1 \times 10^{-10}$ | | | | |
| Pairwise comparisons |  |  |  |  |
| Genotype 1\Genotype 2 | *D. simulans* | *D. mauritiana* | F1 (♀ *sim* × ♂ *mau*) | F1 (♀ *mau* × ♂ *sim*) |
| *D. simulans* | * | < 0.001 | < 0.001 | NA |
| *D. mauritiana* | 42.18 | * | < 0.001 | NA |
| F1 (♀ *sim* × ♂ *mau*) | 9.35 | 45.30 | * | NA |
| F1 (♀ *mau* × ♂ *sim*) | NA | NA | NA | * |

### Perfuming assays

Since F1 individuals are less attractive to pure species than conspecifics, and their CHC profiles are different from their pure species counterparts, we hypothesized that modifying the CHC of the hybrids to be more akin to the profile of the pure species would increase their chance of mating. We also hypothesized that modifying the CHC profile of pure species to resemble the CHC profile of hybrids would decrease their mating success.

First, we studied whether the perfuming treatment affected the CHC profile of perfumed flies. We focused on the *simulans/mauritiana* species pair. Figure 4 shows a PCA of the CHCs of pure *D. simulans*, pure *D. mauritiana*, and the reciprocal F1s (as shown in Figure 4) but also shows the CHC pattern of perfumed *D. simulans* (Figure 4A), perfumed *D. mauritiana* (Figure 4B), and perfumed F1 females (Figure 4C) in PCA biplots. Tables S14-S16 show the loadings of the PCAs, and Figures S8-S10 show the eigenvectors. In the three perfuming experiments, we observed that perfumed individuals had a CHC profile that was intermediate between their genotype and the population to which they were exposed during the perfuming phase (Figure 4, Table 3). These results suggest that perfuming treatments are an effective way to modify, but not fully replace, the CHC of a focal fly.

**TABLE 3.** Perfuming experiments induce differences in the joint CHC profile in *D. simulans, D. mauritiana* and F1 hybrids. We focus on the species of the *D. simulans* species group (*D. simulans* and *D. mauritiana*). We performed pairwise comparisons using a Tukey test following a One-way ANOVA. The lower triangular matrix shows the t value from multiple comparisons of means using Tukey contrasts. The upper triangular matrix shows the P-value associated to the comparison. All P-values were adjusted for multiple comparisons. Please note that we limited our analyses to PC1, because that PC explains over 75% of the variance in both species pairs. Figure 4 shows a representation of the same results.

| ***D. simulans* perfuming experiments: $F_{4,40} = 32.695$, $P < 1 \times 10^{-10}$** |  |  |  |  |  |
| --- | --- | --- | --- | --- | --- |
|  | **Pairwise comparisons** |  |  |  |  |
| **Genotype 1\Genotype 2** | ***D. simulans*** | **F1 (♀ *sim* × ♂ *mau*)** | ***D. simulans* perfumed with F1 (♀ *sim* × ♂ *mau*)** | ***D. simulans* perfumed with *mau*** | ***D. mauritiana*** |
| ***D. simulans*** | * | 0.709 | 1.000 | < 0.001 | < 0.001 |
| **F1 (♀ *sim* × ♂ *mau*)** | 1.267 | * | 0.821 | < 0.001 | < 0.001 |
| ***D. simulans* perfumed with F1 (♀ *sim* × ♂ *mau*)** | 0.147 | 1.064 | * | < 0.001 | < 0.001 |
| ***D. simulans* perfumed with *mau*** | 8.020 | 7.941 | 7.445 | * | 1.000 |
| ***D. mauritiana*** | 6.368 | 6.712 | 6.111 | 0.001 | * |
| ***D. mauritiana* perfuming experiments: $F_{4,43} = 12.584$, $P = 7.305 \times 10^{-7}$** |  |  |  |  |  |

|  | Pairwise comparisons |  |  |  |  |
| --- | --- | --- | --- | --- | --- |
| Genotype 1\Genotype 2 | *D. mauritiana* | F1 (♀ *sim* × ♂ *mau*) | *D. mauritiana* perfumed with F1 (♀ *sim* × ♂ *mau*) | *D. mauritiana* perfumed with *sim* | *D. simulans* |
| *D. mauritiana* | * | < 0.001 | 0.007 | < 0.001 | < 0.001 |
| F1 (♀ *sim* × ♂ *mau*) | 5.548 | * | 0.651 | 0.999 | 0.985 |
| *D. mauritiana* perfumed with F1 (♀ *sim* × ♂ *mau*) | 3.611 | 1.352 | * | 0.570 | 0.824 |
| *D. mauritiana* perfumed with *sim* | 6.780 | 1.721 | 1.481 | * | 0.984 |
| *D. simulans* | 5.800 | 6.015 | 1.050 | 0.523 | * |
| F1 perfuming experiments: $F_{4,32} = 13.472$, $P = 1.522 \times 10^{-6}$ | | | | | |
|  | Pairwise comparisons |  |  |  |  |
| Genotype 1\Genotype 2 | F1 | *D. simulans* | *D. mauritiana* | F1 (♀ *sim* × ♂ *mau*) perfumed with *sim* | F1 (♀ *sim* × ♂ *mau*) perfumed with *mau* |
| F1 (♀ *sim* × ♂ *mau*) | * | 0.567 | < 0.001 | 0.999 | 0.002 |
| ***D. simulans*** | 1.500 | * | < 0.001 | 0.768 | 0.020 |
| ***D. mauritiana*** | 5.861 | 5.168 | * | < 0.001 | 0.205 |
| **F1 (♀ *sim* × ♂ *mau*)**<br>**perfumed**<br>**with *sim*** | 0.219 | 1.165 | 5.369 | * | 0.0056 |
| **F1 (♀ *sim* × ♂ *mau*)**<br>**perfumed**<br>**with *mau*** | 4.225 | 3.260 | 2.196 | 3.754 | * |

Next, we perfumed F1 and pure-species females and studied their attractiveness. The effect of perfuming F1 females was strong for all species, as all male-choice assays showed deviations from a 1:1:1 ratio (expected if there was random-choice; Figure 5); the three types of females (i.e., the type of perfuming treatment) showed differences in attractiveness in all assayed F1 genotypes. F1 females that had been perfumed as pure-species females were more attractive to pure species males, as long as the perfuming treatment and the male species matched (Table S17). F1 females that had been perfumed as other F1s showed a level of attractiveness as expected by random choice (i.e., they were chosen 1/3 of the time). Note that the only difference between these F1 females is whether they were perfumed or not, as their genotype is identical. These results indicate that modifying the CHC profile of F1 females changes their chances of being courted by a pure species male. The blend of CHC in hybrids is an important component of their reduced sexual attractiveness to pure species males, which ultimately affects the possibility they might serve as a bridge for gene flow between species.

**FIGURE 5.**
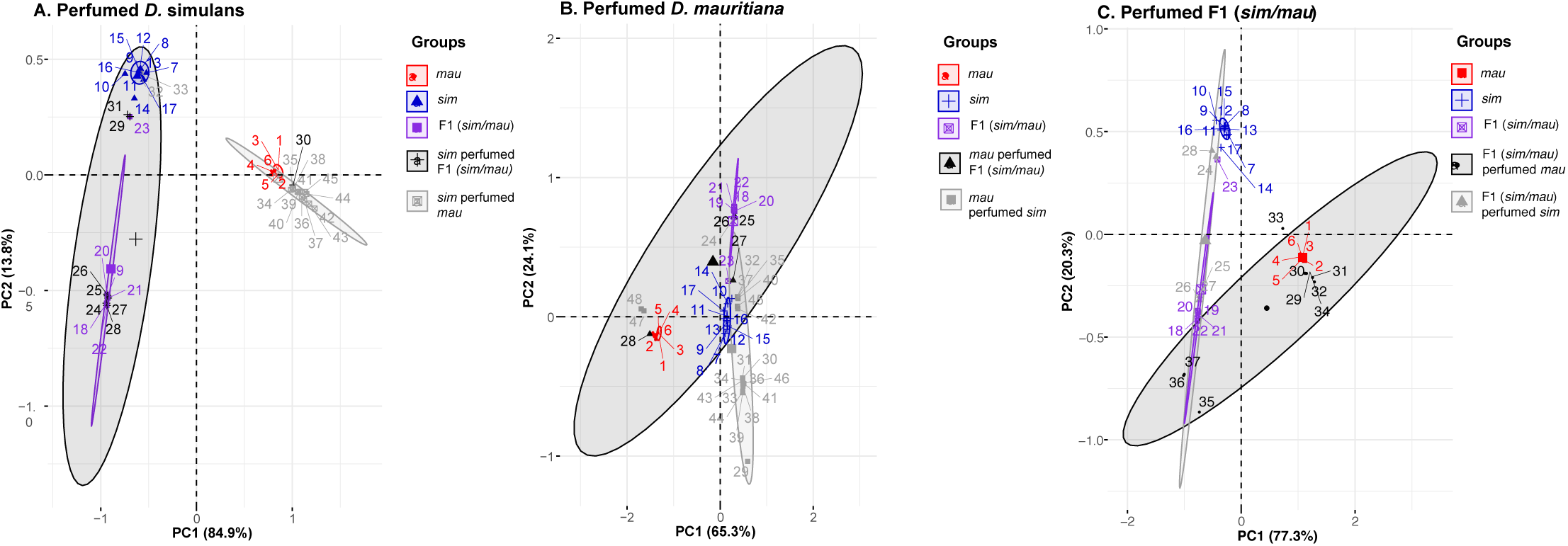
Perfuming hybrid and pure species females from the *simulans* species group modifies their CHC profile. The perfuming treatment consisted of raising a single fly with a group of 10 flies from a different genotype. For this section, we focused on the study of *D. simulans* (A), *D. mauritiana* (B) and F1(♀*sim*/♂*mau*) hybrids (C). Pure species and F1 (*sim*/*mau*) hybrids are shown using the same colors as in Figure 3. (Please note these data is the same as Figure 3). Perfumed samples are shown in one of two shades of gray.) In the legend, the genotype of the fly appears first, and the perfuming treatment (i.e., the genotype of individuals placed along the focal fly) appears second (species/perfume).

Finally, we did choice mating experiments that involved perfuming pure species females. As occurred with the F1 hybrid perfuming experiments, perfuming lead to differences in the pure-species females attractiveness. Even though treatments differ among themselves, no treatment differed from the 1/3 expectation, suggesting relatively mild effects of the perfuming treatment. Females perfumed with the CHCs of their conspecifics were the most attractive type to their conspecific males (Figure 6, Table S18). Pure species females perfumed as heterospecifics showed the lowest level of mating, suggesting that their CHC blend is less attractive to pure species males. Pure species females perfumed with the CHC blend of the hybrids show a decrease compared with pure-species perfumed with their native blend and a level of attractiveness similar to that of pure-species females perfumed with the heterospecific blend. These results are consistent with the possibility that the CHC blend in hybrid females is less attractive to both pure species. The results from our perfuming experiments suggest that hybrid CHC blends are deleterious as they reduce the fitness of pure species individuals that have been perfumed like hybrids.

**FIGURE 6.**
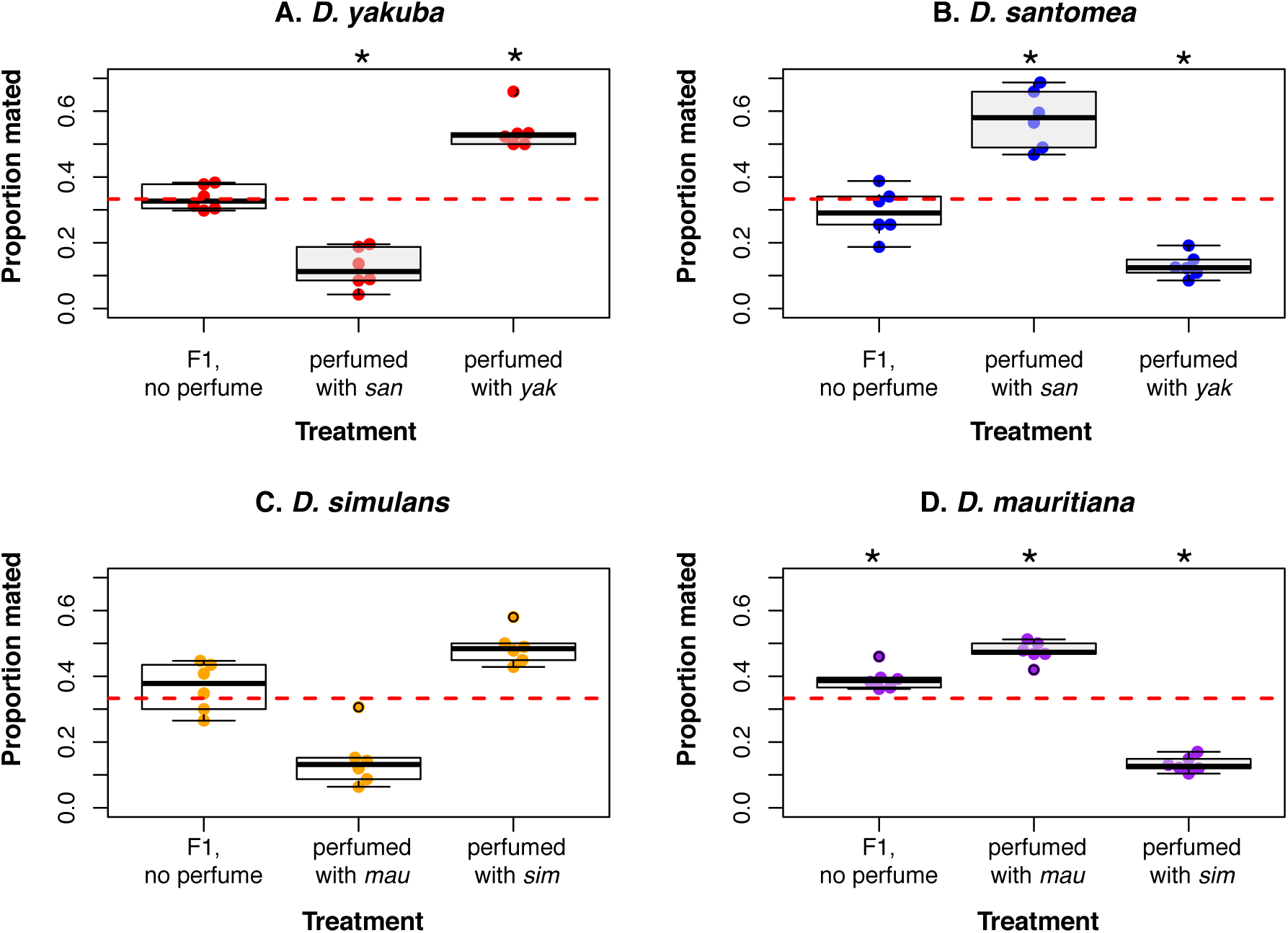
Perfuming hybrid females with the CHC blend of pure species changes their attractiveness to pure species males. Each experiment consisted of a pure species male having the choice of three different females with identical genotypes but differences in their perfuming treatment. Each point shows the proportion of the three types of females chosen in a block of matings (n = 50 observations). The red line shows the expected mating frequencies for the three types of females if perfuming has no effect on sexual attractiveness. Pairwise comparisons between perfuming categories are shown in Table S17. Treatments that significantly differ from the 1/3 expectation are marked with stars.

**FIGURE 7.**
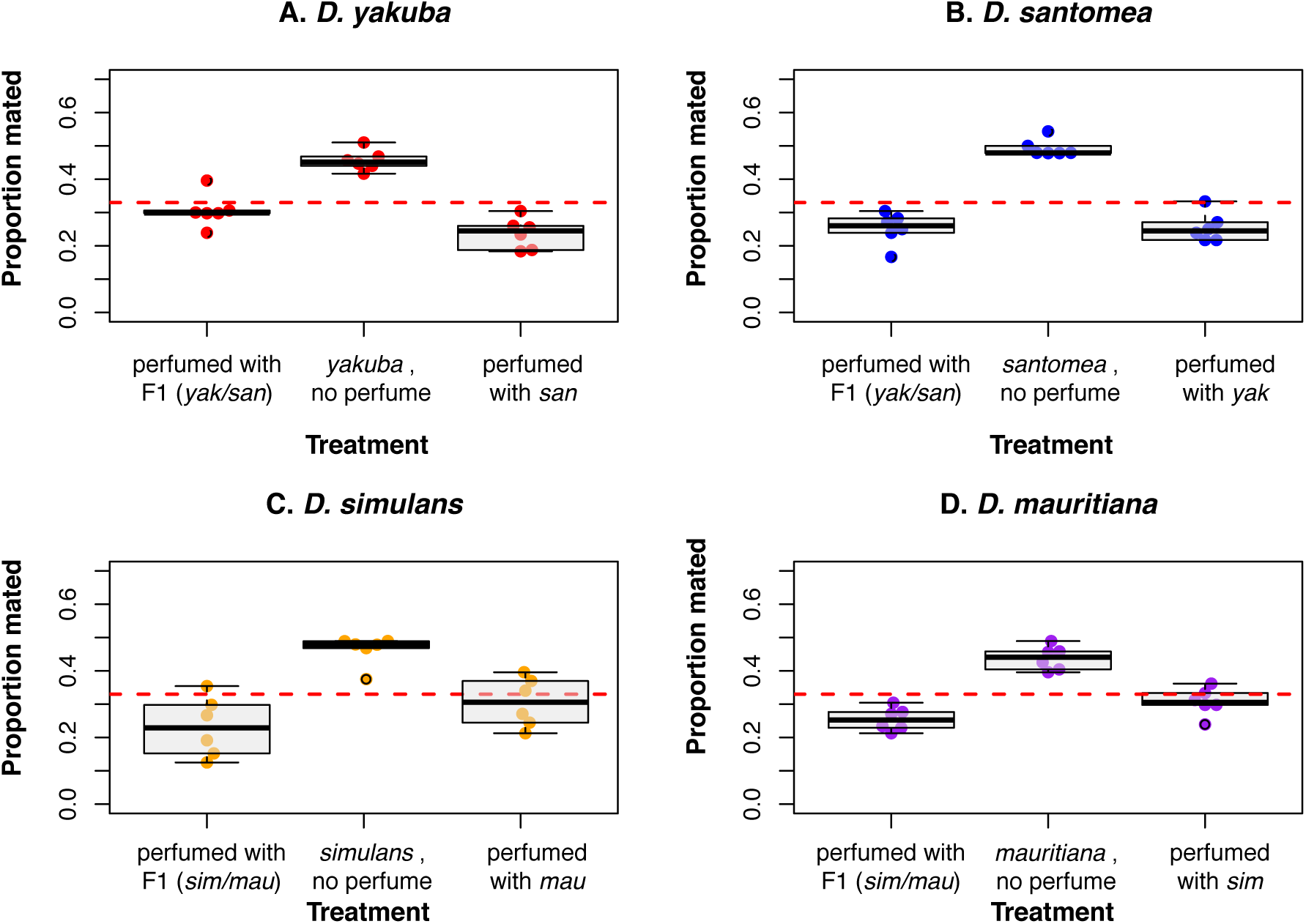
Perfuming pure species females with the CHC blend of heterospecifics or hybrid females reduces their attractiveness to pure species males. Each experiment consisted of a pure species male having the choice of three conspecific females with identical genotypes but differences in their CHC blend. Each point shows the proportion of the three types of females chosen in a block of matings (n = 50 observations). The red line shows the expected mating frequencies for the three types of females if perfuming has no effect on attractiveness. Pairwise comparisons between perfuming categories are shown in Table S18. None of the three treatments significantly differs from the 1/3 expectation.

## DISCUSSION

Prezygotic isolation is common in nature, but the high prevalence of gene flow suggest that prezygotic barriers are leaky (Irwin 2019). Hybridization is a common occurrence in all taxa in which surveys have been systematically performed (Harrison and Larson 2014b; Mallet et al. 2016; Taylor and Larson 2019). Over 10% of animal species hybridize in nature, and the number might be higher for plants and fungi (Schardl and Craven 2003; Mallet 2005; Ellstrand 2014; Mallet et al. 2016). In cases where hybridization occurs, lower hybrid fitness is an important component of how species persist in nature (Coughlan and Matute 2020). In this report, we describe that *Drosophila* hybrids are less attractive to the pure species when the pure-species individuals have a choice. This results indicate that even when hybrids are fertile, they might suffer from subtle defects that reduced their fitness in nature and might limit their ability to serve as genetic bridges for introgression. This defects might not be rare in nature. Insect hybrids show anomalous courtship behavior (Noor 1997; Kost et al. 2016), while salmonid hybrids have trait combinations that make them less attractive to pure species (Fukui et al. 2018).

In the case of the *simulans* and *yakuba* species complex, F1 hybrids have a CHC profile that is intermediate to that of their parents. Discrimination against hybrids seems to be mediated by that intermediate profile. CHCs have been primarily implicated in two important processes in insects: desiccation resistance, and communication (e.g., (Jallon and David 1987; Foley and Telonis-Scott 2011; Arcaz et al. 2016) reviewed in (Gibbs 1998; Chung and Carroll 2015)). *Drosophila* hybrids often show intermediate blends of CHCs (Coyne et al. 1994; Hercus and Hoffmann 1999; Gleason et al. 2009; Combs et al. 2018) which poses the question of how generalizable our results are to other hybrids. At least one case suggests the existence of a similar occurrence in beetles. Species of the beetle genus *Altica* differ in their CHC profile. F1 hybrids between *A. fragariae* and *A. viridicyanea* show CHC profiles that are intermediate from the parents but are also distinct (Xue et al. 2018). Males from the pure species discriminate strongly against hybrid females, potentially cued on their CHC blend profile (Xue et al. 2018). Only a broad-scale survey will reveal whether lower attractiveness is pervasive not only in *Drosophila* but across animals.

CHCs are regularly the target of natural and sexual selection (Menzel et al. 2017); species (Higgie et al. 2000) —and populations (Higgie and Blows 2008; Veltsos et al. 2012)— might differ as a result of local adaptation. There is no strong difference in desiccation resistance between *D. simulans* and *D. mauritiana*; differences between *D. simulans* lines are larger than the differences between species (Van Herrewege and David 1997). *Drosophila santomea* is slightly more resistant than *D. yakuba* to desiccation (Matute and Harris 2013), which might be explained not by its particular CHC blend of the species but by its higher total CHC content in the cuticle (Mas and Jallon 2005). Even though *D. simulans* and *D. yakuba* are human commensals that tend to be found in dryer environments, the ecological effects behind the similarities and differences in CHC profiles in these two species pairs remain unknown.

The two *Drosophila* species pairs studied here exchange alleles in nature. *Drosophila yakuba* and *D. santomea* form a hybrid zone in the midlands of São Tomé, where 3-5% of the collected individuals (both males and females) from the *yakuba* clade are hybrids (Comeault et al. 2016; Turissini and Matute 2017). To date, *D. simulans* and *D. mauritiana* are not known to form an extant hybrid zone. In both cases, species boundaries are porous and have allowed for introgression between species, but the introgression between the two species is less than 1% per genome per individual on average (Kliman et al. 2000; Bachtrog et al. 2006; Turissini and Matute 2017; Meiklejohn et al. 2018). Hybrid males from the two species in this study are sterile, and their fitness is effectively zero (Coyne 1985; Coyne et al. 2004). Hybrid male sterility is a stronger form of isolation than the lower male sexual attractiveness reported here. However, hybrid females from these species are fertile and can interbreed with males from both pure species. The existence of hybrid defects that lead to selection against F1 hybrids might be important in the persistence of species that hybridize in nature.

Our study is limited in that it does not recapitulate the between-sexes interactions that occur in nature. We cannot estimate the full extent of the fitness reduction that the lower sexual attractiveness might cause. Field experiments of paternity and rates of insemination of hybrid females can reveal whether these defects also occur in nature. F1 hybrid stickleback males in natural enclosure experience strong sexual selection against them as evidenced by the observation that limnetic males are more vigorous in their display towards limnetic females —a proxy of mating success— than hybrid males (Vamosi and Schluter 1999).

Comparative analyses have suggested that premating behavioral isolation is completed relatively faster than hybrid sterility and inviability, and thus might play an important role in setting the speciation process in motion (Coyne and Orr 1989, 1997; Sasa et al. 1998; Moyle et al. 2004; Rabosky and Matute 2013; Castillo 2017). Nonetheless, postzygotic isolation plays an important role in keeping species apart and on completing prezygotic isolation via reinforcement (Rosenblum et al. 2012, Coughlan and Matute 2020). Other forms of prezygotic isolation, not related to mating behavior, also seem to evolve quickly (Turelli et al. 2014; Turissini et al. 2017). Future studies should measure the rate of evolution of behavioral postzygotic isolation and assess whether it is more akin to the rate of evolution of premating isolation or to that of hybrid inviability and sterility. They should also compare the magnitude of the hybrid defect in homo- and heterogametic sexes which would reveal whether Haldane’s rule occurs in behavioral postzygotic isolation.

Our focus on this study was to assess whether *Drosophila* hybrids suffer from reduced attractiveness. Hybrid fitness is a continuum that ranges from hybrid vigor to complete inviability (Guerrero et al. 2017; Dagilis et al. 2019). In some cases, hybrids might be more attractive to their parentals that their own conspecifics (Pfennig 2007). Hybrids might also be less attractive to the pure species but more attractive to other hybrids thus facilitating hybrid speciation (e.g., (Mavárez et. al 2006; Melo et al. 2009; Selz et al. 2014; Schmidt and Pfennig 2016; Comeault and Matute 2018). Only a concerted effort to dissect the multiple fitness components of hybrids will reveal whether discrimination against hybrids is widespread in nature and important for species persistence, or on the contrary is an barrier to gene flow exclusively restricted to *Drosophila*.

## ACKNOWLEDGMENTS

We would like to thank A.A. Comeault, B.S. Cooper and the members of the Matute lab for helpful scientific discussions and comments. We would like to thank the Bioko Biodiversity Protection Program (Equatorial Guinea), and the Ministry of Environment (Republic of São Tomé and Príncipe) for permission to collect and export specimens for study. D.R.M. was supported by the National Science Foundation (Award 1737752). The authors have no conflicts of interest.

