## Supplementary material for "Pure species discriminate against hybrids in the *Drosophila melanogaster* species subgroup": Tables S1-S18, Figures S1-S10

**TABLE OF CONTENTS**

**TABLE S1. Cuticular hydrocarbons (CHCs) included in this study…………………3**
**TABLE S2. Number of samples per genotype used to score CHC profiles………...3**
**TABLE S3. No effect of marking females with colored food in the outcome of the mating in mass matings……………..………………………………………………4**
**TABLE S4. No effect of marking females with wing clipping in the outcome of the mating in mass matings…………………………………………………………......4**
**TABLE S5. No effect of the dual marking scheme in the outcome of the mating in mass matings. ……………………………………………………………………………4**
**TABLE S6. No effect of the dual marking scheme in female attractiveness………..5**
**TABLE S7. Time spent by pure species males courting four different types of females in choice experiments. ……………………………………………………6**
**TABLE S8. Male genotype has a strong effect on the outcome of the mating in non-choice matings………………………………………………………………...7**
**TABLE S9. Pure species females discriminate against heterospecific and F1 hybrid males in no-choice mating trials…………………………………..……8**
**TABLE S10. Male effort in matings with conspecific, heterospecific, and hybrid males in no-choice experiments…………………………………………………..10**
**TABLE S11. Copulation duration in matings with conspecific, heterospecific, and hybrid males in no-choice experiments……………………………………….13**
**TABLE S12. PC loadings for individual CHC peaks based on PCA for pure species and F1s in the *D. yakuba/D. santomea* species pair…………………………..14**
**TABLE S13. PC loadings for individual CHC peaks based on PCA for pure species and F1s in the *D. simulans/D. mauritiana* species pair………………………15**
**TABLE S14. PC loadings for individual CHC peaks based on PCA for D. *simulans*, *D. mauritiana*, F1 (*sim/mau*) and perfumed *D. simulans* samples…………15**
**TABLE S15. PC loadings for individual CHC peaks based on PCA for D. *simulans*, *D. mauritiana*, F1 (*sim/mau*) and perfumed *D. mauritiana* samples……….15**

**FIGURE S3. No effect of the marking scheme. Time spent courting each type of female. ……………………………………………………………………………..24**
**FIGURE S4. Regression curves for seven CHCs. Each panel shows one CHC… ……………………………………………………………………………………..25**
**FIGURE S5. No strong effect of the scorer on the proxy of female attractiveness. ……………………………………………………………………………………..26**
**FIGURE S6. PC1 and PC2 eigenvectors for the *D. yakuba/D. santomea* biplot. ……………………………………………………………………………………..27**

**FIGURE S7. PC1 and PC2 eigenvectors for the *D. simulans/D. mauritiana* biplot. ……………………………………………………………………………………..28**
**FIGURE S8. PC1 and PC2 eigenvectors for the perfumed *D. simulans* samples biplot……………………………………………………………………………..29**
**FIGURE S9. PC1 and PC2 eigenvectors for the perfumed *D. mauritiana* samples biplot……………………………………………………………………………...30**
**FIGURE S10. PC1 and PC2 eigenvectors for the perfumed F1 *(sim/mau)* samples biplot. ……………………………………………………………………………31**

### SUPPLEMENTARY TABLES

**TABLE S1. Cuticular hydrocarbons (CHCs) included in this study. The standard curves were derived from serial dilutions.** The regression curve for each compound is show in Figure S4.

| Compound | Catalog number | Elution time | Standard curve |
|---|---|---|---|
| n-Heneicosane | Sigma Aldrich, 51523-1g | 14.482-14.56 | y=2.248*x |
| 11-*cis*-Vaccenyl Acetate | Cayman chemical, #10010101 | 17.389-17.42 | y=1.189*x |
| 7(Z)-tricosene | Cayman chemical, #9000313 | 18.64-18.83 | y=1.249*x |
| Tricosane | Sigma Aldrich, 263850 | 23.6-23.68 | y=0.296*x |
| 7-11-pentacosene | Cayman chemical, #9000530 | 26.67-26.856 | y=1.017*x |
| 7(Z),11(Z)-Heptacosadiene | Cayman chemical, #10012567 | 31.216-31.76 | y=1.586*x |
| 7(Z)-Nonacosadiene | Cayman chemical, #9000314 | 36.7-37.189 | y=1.095*x |

**TABLE S2. Number of samples per genotype used to score CHC profiles.**

| Genotype | N |
|---|---|
| *D. yakuba* | 13 |
| *D. santomea* | 15 |
| F1 (♀*yak*/♂*san*) | 8 |
| F1 (♀*san*/♂*yak*) | 21 |
| *D. simulans* | 6 |
| *D. mauritiana* | 11 |
| F1 (♀*sim*/♂*mau*) | 6 |
| *D. simulans* perfumed with F1 (♀sim/♂mau) | 8 |
| *D. simulans* perfumed with *D. mauritiana* | 14 |
| *D. mauritiana* perfumed with F1 | 4 |
| *D. mauritiana* perfumed with *D. simulans* | 21 |
| F1 perfumed with *D. simulans* | 5 |
| F1 perfumed with *D. mauritiana* | 9 |

**TABLE S3. No effect of marking females with colored food in the outcome of the mating in mass matings.** We scored 100 females for each species. Mated flies shows the total number of mated males. df: degrees of freedom.

| Species | N | $X^2$ | Df | P |
|---|---|---|---|---|
| *D. yakuba* | 100 | 1.781 | 3 | 0.619 |
| *D. santomea* | 100 | 5.156 | 3 | 0.161 |
| *D. simulans* | 100 | 1.026 | 3 | 0.795 |
| *D. mauritiana* | 100 | 2.240 | 3 | 0.524 |

**TABLE S4. No effect of marking females with wing clipping in the outcome of the mating in mass matings.** We scored 100 females for each species. Mated flies shows the total number of mated males. df: degrees of freedom.

| Species | N | $X^2$ | df | P |
|---|---|---|---|---|
| *D. yakuba* | 100 | 1.010 | 3 | 0.799 |
| *D. santomea* | 100 | 3.297 | 3 | 0.348 |
| *D. simulans* | 100 | 1.371 | 3 | 0.712 |
| *D. mauritiana* | 100 | 1.663 | 3 | 1.663 |

**TABLE S5. No effect of the dual marking scheme in the outcome of the mating in mass matings. Flies were labeled with abdominal colors and by clipping their wings.** We scored 480 females for each species. Mated flies shows the total number of mated males. df: degrees of freedom.

| Species | Mated flies | $X^2$ | df | P |
|---|---|---|---|---|
| ***D. yakuba*** | 428 | 5.606 | 15 | > 0.9 |
| ***D. santomea*** | 387 | 6.096 | 15 | > 0.9 |
| ***D. simulans*** | 402 | 2.263 | 15 | > 0.9 |
| ***D. mauritiana*** | 422 | 4.069 | 15 | > 0.9 |

**TABLE S6. No effect of the dual marking scheme in female attractiveness in mating choices where males were given the choice between four conspecific females and different markings.** The proxy for attractiveness is the time spent by the male courting each type of female. None of the dual markings, as determined by the interaction between effects, had a strong effect on the time that a male spent courting different types of females.

| **Species** | **Effect** | **$F_{9,80}$** | **P** |
|---|---|---|---|
| *D. yakuba* | Color | 0.643 | 0.590 |
| *D. yakuba* | Clipping | 0.114 | 0.952 |
| *D. yakuba* | Color × clipping | 0.560 | 0.825 |
| *D. santomea* | Color | 0.677 | 0.569 |
| ***D. santomea*** | **Clipping** | **2.776** | **0.047** |
| *D. santomea* | Color × clipping | 1.665 | 0.111 |
| *D. simulans* | Color | 0.834 | 0.479 |
| *D. simulans* | Clipping | 0.767 | 0.516 |
| *D. simulans* | Color × clipping | 1.242 | 0.282 |
| *D. mauritiana* | Color | 0.339 | 0.797 |
| *D. mauritiana* | Clipping | 2.263 | 0.087 |
| *D. mauritiana* | Color × clipping | 0.575 | 0.814 |

**TABLE S7. Time spent by pure species males courting four different types of females in choice experiments.** Means show the time a male spent courting each type of female relative to the total time spent courting all females . All the means and standard deviations (SD) are based on 300 observations. The last four columns show pairwise comparisons as 4 × 4 matrices for each cross. The lower triangular matrix shows the t value from a multiple comparisons of means using Tukey contrasts. The upper triangular matrix shows the P-value associated to the comparison. All p-values were adjusted for multiple comparisons.

| ***D. yakuba* males:** LMM, $F_{3,957}$ = 9,133.095 P <0.0001 | | | | | |
|---|---|---|---|---|---|
| | | Pairwise comparisons | | | |
| Female genotype | Mean (SD) | *D. yakuba* | *D. santomea* | F1 (♀*yak* × ♂*san*) | F1 (♀*san* × ♂*yak*) |
| *D. yakuba* | 0.744 (0.080) | * | <0.001 | <0.001 | <0.001 |
| D. santomea | 0.107 (0.065) | 130.569 | * | <0.001 | <0.001 |
| F1 (♀*yak* × ♂*san*) | 0.073 (0.030) | 137.538 | 6.968 | * | 0.923 |
| F1 (♀*san* × ♂*yak*) | 0.076 (0.053) | 136.911 | 6.341 | -0.627 | * |
| ***D. santomea* males:** LMM, $F_{3,957}$ = 6,021.967 <0.0001 | | | | | |
| | | Pairwise comparisons | | | |
| Female genotype | Mean (SE) | *D. yakuba* | *D. santomea* | F1 (♀*yak* × ♂*san*) | F1 (♀*san* × ♂*yak*) |
| *D. santomea* | 0.704 (0.086) | * | <0.001 | <0.001 | <0.001 |
| *D. yakuba* | 0.089 (0.004) | 111.260 | * | 0.222 | <0.001 |
| F1 (♀*yak* × ♂*san*) | 0.129 (0.072) | 7.249 | 103.981 | * | <0.001 |
| F1 (♀*san* × ♂*yak*) | 0.079 (0.048) | 1.917 | 113.144 | 9.160 | * |
| ***D. simulans* males:** LMM, $F_{3,957}$: 10,462.75 <0.0001 | | | | | |
| | | Pairwise comparisons | | | |
| Female genotype | Mean (SD) | *D. simulans* | *D. mauritiana* | F1 (♀*sim* × ♂*mau*) | F1 (♀*mau* × ♂*sim*) |

| | | | | | |
|---|---|---|---|---|---|
| *D. simulans* | 0.760 (0.074) | * | <0.001 | <0.001 | <0.001 |
| *D. mauritiana* | 0.111 (0.058) | 137.738 | * | <0.001 | <0.001 |
| F1 (♀*sim* × ♂*mau*) | 0.071 (0.052) | 146.178 | 8.458 | * | 0.035 |
| F1 (♀*mau* × ♂*sim*) | 0.058 (0.042) | 148.977 | 11.140 | 2.709 | * |
| ***D. mauritiana* males,** LMM: $F_{3,957}$: 25,822.28, P <0.0001 | | | | | |
| | | Pairwise comparisons | | | |
| | Mean (SD) | *D. mauritiana* | *D. simulans* | F1 (♀*sim* × *mau*) | F1 (♀*mau* × *sim*) |
| *D. mauritiana* | 0.867 (0.061) | * | <0.001 | <0.001 | <0.001 |
| *D. simulans* | 0.080 (0.057) | 217.242 | * | <0.001 | <0.001 |
| F1 (♀*sim* × ♂*mau*) | 0.036 (0.027) | 229.029 | 11.922 | * | <0.001 |
| F1 (♀*mau* × ♂*sim*) | 0.017 (0.013) | 234.376 | 17.292 | 5.413 | * |

**TABLE S8. Male genotype has a strong effect on the outcome of the mating in non-choice matings.** We fit a linear mixed model in which the only fixed effect was the genotype of the male. The response was whether a female mated or not (scored as a binomial outcome). The significance of  effect was calculated using likelihood ratio tests. Df: degrees of freedom, LRT: Likelihood ratio test

| **Female** | **df** | **LRT** | **P** |
|---|---|---|---|
| *D. yakuba* | 3 | 1,747.8 | $< 1 \times 10^{-10}$ |
| *D. santomea* | 3 | 3,280.9 | $< 1 \times 10^{-10}$ |
| *D. simulans* | 3 | 2,487.3 | $< 1 \times 10^{-10}$ |
| *D. mauritiana* | 3 | 3,227.0 | $< 1 \times 10^{-10}$ |

**TABLE S9. Pure species females discriminate against heterospecific and F1 hybrid males in no-choice mating trials.** We used a logistic regression to calculate the likelihood of mating between pure species females and males from four different genotypes. We used a Tukey test to do pairwise comparisons between male genotypes. The last four columns show pairwise comparisons as 4 × 4 matrices for each cross. The lower triangular matrix shows the t value from a multiple comparisons of means using Tukey contrasts. The upper triangular matrix shows the P-value associated to the comparison. Please note that results are given on the log odds ratio (not the response) scale. yak: *D. yakuba*; *san: D. santomea.*

| **Female** | **Male** | **Least squares mean (SE)** | **Pairwise comparisons** | | | |
|---|---|---|---|---|---|---|
| | | | *yak* | *san* | F1 (♀*yak* × ♂*san*) | F1 (♀*san* × ♂*yak*) |
| *yak* | *yak* | 2.782 (0.168) | * | <.0001 | <.0001 | <.0001 |
| | *san* | 1.141 (0.126) | 25.540 | * | 0.377 | 0.442 |
| | F1 (♀*yak* × ♂*san*) | -1.312 (0.128) | 26.356 | 1.602 | * | 0.999 |
| | F1 (♀*san* × ♂*yak*) | -1.300 (0.128) | -26.301 | -1.493 | 0.109 | * |
| **Female** | **Male** | **Least squares mean (SE)** | **Pairwise comparisons** | | | |
| | | | *san* | *yak* | F1 (♀*yak* × ♂*san*) | F1 (♀*san* × ♂*yak*) |
| *san* | *san* | 2.719 (0.237) | * | <.0001 | <.0001 | <.0001 |
| | *yak* | -3.220 (0.253) | 27.031 | * | 0.0015 | 0.073 |

| | | | | | | |
|---|---|---|---|---|---|---|
| | F1 (♀*yak* × ♂*san*) | -4.294 (0.323) | 23.532 | -3.643 | * | 0.488 |
| | F1 (♀*san* × ♂*yak*) | -3.836 (0.286) | 25.460 | -2.422 | 1.418 | * |
| **Female** | **Male** | **Least squares mean (SE)** | **Pairwise comparisons** | | | |
| | | | *sim* | *mau* | F1 (♀*sim* × ♂*mau*) | F1 (♀*mau* × ♂*sim*) |
| *sim* | *sim* | 2.649 (0.164) | * | <.0001 | <.0001 | <.0001 |
| | *mau* | -1.517 (0.133) | -27.302 | * | <.0001 | <.0001 |
| | F1 (♀*sim* × ♂*mau*) | -2.558 (0.160) | -29.443 | 7.149 | * | 1.0000 |
| | F1 (♀*mau* × ♂*sim*) | -2.558 (0.160) | -29.445 | 7.149 | 0.000 | * |
| **Female** | **Male** | **Least squares mean (SE)** | **Pairwise comparisons** | | | |
| | | | *mau* | *sim* | F1 (♀*sim* × ♂*mau*) | F1 (♀*mau* × ♂*sim*) |
| *mau* | *mau* | 2.154 (0.139) | * | <.0001 | <.0001 | <.0001 |
| | *sim* | -4.541 (0.319) | 20.584 | * | 0.9315 | 0.566 |

| | | | | | | |
|---|---|---|---|---|---|---|
| | F1 (♀*sim* × ♂*mau*) | -4.296 (0.286) | 21.966 | 0.602 | * | 0.893 |
| | F1 (♀*mau* × ♂*sim*) | -4.041 (0.257) | -6.194 | 1.296 | 0.711 | * |

**TABLE S10. Male effort in matings with conspecific, heterospecific, and hybrid males in no-choice experiments.** Effort is defined as the number of time windows (measured every 2 minutes) that a male courted the female divided by time to mating (latency). Matings with heterospecific and hybrid males take longer to occur than conspecific matings. *N* represents the number of mated pairs used for the analyses. All the means (percentage of females mated) and standard deviations (SD). The last four columns show pairwise comparisons as 4 × 4 matrices for each cross. The upper triangular matrix shows the t value from a multiple comparisons of means using Tukey contrasts. The lower triangular matrix shows the P-value associated to the comparison. All p-values were adjusted for multiple comparisons.

| ***D. yakuba* females**: LMM, $F_{3,1606}$ = 125.98 < 0.00001 | | | | | | |
|---|---|---|---|---|---|---|
| Male genotype | N | Mean (SD) | Pairwise comparisons | | | |
| | | | *D. yakuba* | *D. santomea* | F1 (♀*yak* × ♂*san*) | F1 (♀*san* × ♂*yak*) |
| *D. yakuba* | 939 | 0.259 (0.112) | * | <.0001 | <.0001 | <.0001 |
| *D. santomea* | 247 | 0.159 (0.100) | 13.298 | * | 0.899 | 0.394 |
| F1 (♀*yak* × ♂*san*) | 214 | 0.166 (0.092) | 11.709 | 0.685 | * | 0.129 |
| F1 (♀*san* × ♂*yak*) | 210 | 0.144 (0.084) | 14.373 | 1.558 | 2.165 | * |
| ***D. santomea* females:** LMM, $F_{3,1009}$ = 4.2138, P = 0.005667 | | | | | | |
| Male genotype | N | Mean (SD) | Pairwise comparisons | | | |
| | | | *D. santomea* | *D. yakuba* | F1 (♀*yak* × ♂*san*) | F1 (♀*san* × ♂*yak*) |
| *D. santomea* | 928 | 0.182 (0.105) | * | 0.057 | 0.525 | 0.085 |
| *D. yakuba* | 45 | 0.143 (0.068) | 2.481 | * | 0.981 | 0.999 |

| | | | | | | |
|---|---|---|---|---|---|---|
| F1 (♀*yak* × ♂*san*) | 15 | 0.147 (0.083) | 1.320 | 0.371 | * | 0.977 |
| F1 (♀*san* × ♂*yak*) | 25 | 0.134 (0.064) | 2.326 | 0.118 | 0.391 | * |
| ***D. simulans* females:** LMM, $F_{3,1262} = 79.441$, $P < 0.0001$ | | | | | | |
| Male genotype | N | Mean (SD) | Pairwise comparisons | | | |
| | | | *D. simulans* | *D. mauritiana* | F1 (♀*sim* × ♂*mau*) | F1 (♀*mau* × ♂*sim*) |
| *D. simulans* | 931 | 0.176 (0.106) | * | <0.001 | <0.001 | <0.001 |
| *D. mauritiana* | 185 | 0.079 (0.066) | 12.362 | * | 0.442 | 0.832 |
| F1 (♀*sim* × ♂*mau*) | 75 | 0.099 (0.087) | 6.622 | 1.463 | * | 0.207 |
| F1 (♀*mau* × ♂*sim*) | 75 | 0.068 (0.040) | 9.233 | 0.826 | 1.919 | * |
| ***D. mauritiana* females:** LMM, $F_{3,929} = 7.5084$, $P < 0.00001$ | | | | | | |
| Male genotype | N | Mean (SD) | Pairwise comparisons | | | |
| | | | *D. mauritiana* | *D. simulans* | F1 (♀*sim* × ♂*mau*) | F1 (♀*mau* × ♂*sim*) |
| *D. mauritiana* | 894 | 0.179 (0.117) | * | 0.0120 | 0.063 | 0.023 |
| *D. simulans* | 11 | 0.072 (0.051) | 3.035 | * | 0.909 | 0.976 |
| F1 (♀*sim* × ♂*mau*) | 14 | 0.102 (0.116) | 2.453 | 0.645 | * | 0.999 |
| F1 (♀*mau* × ♂*sim*) | 14 | 0.097 (0.045) | 2.816 | 0.403 | 0.258 | * |

**TABLE S11. Copulation duration in matings with conspecific, heterospecific, and hybrid males in no-choice experiments.** Matings with heterospecific and hybrid males take longer to occur than conspecific matings. *N* represents the number of mated pairs used for the analyses. All the means (percentage of females mated) and standard deviations (SD). The last four columns show pairwise comparisons as 4 × 4 matrices for each cross. The upper triangular matrix shows the t value from a multiple comparisons of means using Tukey contrasts. The lower triangular matrix shows the P-value associated to the comparison. All p-values were adjusted for multiple comparisons.

| ***D. yakuba* females:** LMM, $F_{3, 1606}$ = 160.89, P < 0.00001 | | | | | | |
|---|---|---|---|---|---|---|
| Male genotype | N | Mean (SD) | Pairwise comparisons | | | |
| | | | *yak* | *san* | F1 (♀*yak* × ♂*san*) | F1 (♀*san* × ♂*yak*) |
| *yak* | 939 | 25.693 (3.888) | * | <.0001 | <.0001 | 0.654 |
| *san* | 247 | 19.343 (3.618) | 20.530 | * | <.0001 | <.0001 |
| F1 (♀*yak* × ♂*san*) | 214 | 22.208 (6.383) | 10.636 | 7.093 | * | <.0001 |
| F1 (♀*san* × ♂*yak*) | 210 | 25.315 (4.355) | 1.144 | 14.710 | 7.396 | * |
| ***D. santomea* females:** LMM, $F_{3,1009}$ = 9.8807, P < 0.00001 | | | | | | |
| Male genotype | N | Mean (SD) | Pairwise comparisons | | | |
| | | | *san* | *Yak* | F1 (♀*yak* × ♂*san*) | F1 (♀*san* × ♂*yak*) |
| *san* | 928 | 29.691 (8.035) | * | < 0.001 | 0.120 | 0.007 |
| *yak* | 45 | 24.708 (5.653) | 4.050 | * | 0.998 | 0.999 |
| F1 (♀*yak* × ♂*san*) | 15 | 25.138 (11.694) | 2.170 | 0.179 | * | 0.994 |
| F1 (♀*san* × ♂*yak*) | 25 | 24.498 (10.029) | 3.178 | 0.104 | 0.243 | * |
| ***D. simulans* females:** LMM, $F_{3,1262}$ = 311.49, P < 0.00001 | | | | | | |

| Male genotype | N | Mean (SD) | Pairwise comparisons | | | |
|---|---|---|---|---|---|---|
| | | | *sim* | *Mau* | F1 (♀*sim* × ♂*mau*) | F1 (♀*mau* × ♂*sim*) |
| *sim* | 931 | 31.981 (5.967) | * | <0.001 | <0.001 | <0.001 |
| *mau* | 185 | 18.126 (8.149) | 28.276 | * | <0.001 | <0.001 |
| F1 (♀*sim* × ♂*mau*) | 75 | 22.493 (3.281) | 12.985 | 5.241 | * | 0.0806 |
| F1 (♀*mau* × ♂*sim*) | 75 | 24.831 (2.924) | 9.786 | 8.047 | 2.352 | * |
| ***D. mauritiana* females:** LMM, $F_{3,929}$ = 25.026, P < 0.00001 | | | | | | |
| Male genotype | N | Mean (SD) | Pairwise comparisons | | | |
| | | | *mau* | *Sim* | F1 (♀*sim* × ♂*mau*) | F1 (♀*mau* × ♂*sim*) |
| *mau* | 894 | 29.373(6.799) | * | < 0.001 | < 0.001 | 0.00171 |
| *sim* | 11 | 16.891 (6.298) | 6.056 | * | 0.689 | 0.131 |
| F1 (♀*sim* × ♂*mau*) | 14 | 19.816 (5.064) | 5.222 | 1.069 | * | 0.644 |
| F1 (♀*mau* × ♂*sim*) | 14 | 22.741 (8.213) | 3.624 | 2.137 | 1.139 | * |

**TABLE S12. PC loadings for individual CHC peaks based on PCA for pure species and F1s in the *D. yakuba/D. santomea* species pair.**

| Compound | PC1 | PC2 | PC3 | PC4 | PC5 | PC6 | PC7 |
|---|---|---|---|---|---|---|---|
| Heneicosane | 0.0074 | -0.907 | -0.419 | 0.014 | 0.027 | -0.029 | -0.003 |
| Vaccenyl acetate | 0.005 | -0.108 | 0.1903 | 0.035 | -0.952 | -0.210 | 0.007 |
| Heptacosadiene | 0.0160 | -0.351 | 0.736 | -0.527 | 0.123 | 0.205 | 0.009 |
| 7-11-pentacosene | 0.004 | -0.011 | 0.099 | -0.171 | 0.225 | -0.953 | -0.034 |
| Z-7-tricosene | 0.999 | 0.017 | -0.0193 | -0.008 | -0.002 | 0.002 | 0.002 |
| Tricosane | 0.020 | -0.204 | 0.486 | 0.831 | 0.1635 | -0.057 | -0.018 |
| Nonacosadiene | -0.002 | -0.003 | 0.003 | 0.014 | 0.0168 | -0.034 | 0.999 |

**TABLE S13. PC loadings for individual CHC peaks based on PCA for pure species and F1s in the *D. simulans/D. mauritiana* species pair.**

| | PC1 | PC2 | PC3 | PC4 | PC5 | PC6 | PC7 |
|---|---|---|---|---|---|---|---|
| Heneicosane | -0.004 | 0.106 | 0.474 | -0.238 | 0.362 | -0.760 | 0 |
| Vaccenyl acetate | 0.086 | -0.066 | 0.177 | -0.910 | -0.231 | 0.275 | 0 |
| Heptacosadiene | -0.119 | 0.182 | 0.759 | 0.301 | -0.510 | 0.162 | 0 |
| 7-11-pentacosene | 0.298 | -0.913 | 0.241 | 0.131 | 0.041 | 0.0005 | 0 |
| Z-7-tricosene | -0.940 | -0.327 | -0.031 | -0.082 | -0.002 | -0.035 | 0 |
| Tricosane | -0.072 | 0.103 | 0.331 | 0.032 | 0.744 | 0.565 | 0 |
| Nonacosadiene | 0 | 0 | 0 | 0 | 0 | 0 | 1 |

**TABLE S14. PC loadings for individual CHC peaks based on PCA for D*. simulans*, *D mauritiana*, F1 (*sim/mau*) and perfumed *D. simulans* samples.**

| | PC1 | PC2 | PC3 | PC4 | PC5 | PC6 | PC7 |
|---|---|---|---|---|---|---|---|
| Heneicosane | -0.001 | 0.088 | -0.329 | 0.182 | -0.514 | -0.766 | 0 |
| Vaccenyl acetate | -0.064 | -0.071 | 0.023 | -0.695 | -0.662 | 0.262 | 0 |
| Heptacosadiene | 0.097 | 0.182 | -0.851 | -0.341 | 0.332 | 0.083 | 0 |
| 7-11-pentacosene | -0.277 | -0.923 | -0.250 | 0.081 | 0.032 | -0.0003 | 0 |
| Z-7-tricosene | 0.953 | -0.296 | 0.026 | -0.009 | -0.055 | -0.012 | 0 |
| Tricosane | 0.033 | 0.115 | -0.322 | 0.601 | -0.427 | 0.581 | 0 |
| Nonacosadiene | 0 | 0 | 0 | 0 | 0 | 0 | 1 |

**TABLE S15. PC loadings for individual CHC peaks based on PCA for D*. simulans*, *D. mauritiana*, F1 (*sim/mau*) and perfumed *D. mauritiana* samples.**

| | PC1 | PC2 | PC3 | PC4 | PC5 | PC6 | PC7 |
|---|---|---|---|---|---|---|---|
| Heneicosane | 0.068 | -0.158 | -0.067 | -0.652 | -0.574 | -0.460 | 1.13E-03 |
| Vaccenyl acetate | 0.119 | 0.014 | 0.076 | -0.673 | 0.208 | 0.696 | -2.38E-03 |
| Heptacosadiene | 0.149 | -0.779 | 0.597 | 0.116 | 0.0008 | 0.0370 | -2.01E-04 |
| 7-11-pentacosene | 0.145 | 0.605 | 0.760 | -0.005 | -0.174 | -0.071 | -7.68E-04 |
| Z-7-tricosene | -0.965 | -0.037 | 0.218 | -0.136 | 0.022 | 0.0040 | -4.51E-05 |
| Tricosane | -0.079 | -0.0177 | -0.095 | 0.301 | -0.772 | 0.546 | -3.90E-04 |
| Nonacosadiene | 0.0003 | 0.0005 | 0.0009 | -0.0007 | 0.0007 | 0.002 | 1.00E+00 |

**TABLE S16. PC loadings for individual CHC peaks based on PCA for *D. simulans*, *D mauritiana*, F1 (*sim/mau*) and perfumed F1 (*sim/mau*) samples.**

| | PC1 | PC2 | PC3 | PC4 | PC5 | PC6 | PC7 |
|---|---|---|---|---|---|---|---|
| Heneicosane | -0.002 | 0.060 | -0.371 | 0.020 | -0.372 | 0.181 | -0.829 |
| Vaccenyl acetate | -0.117 | -0.103 | -0.140 | -0.840 | -0.308 | -0.383 | 0.090 |
| Heptacosadiene | 0.121 | 0.114 | -0.503 | 0.386 | -0.572 | -0.163 | 0.465 |
| 7-11-pentacosene | -0.401 | -0.886 | -0.071 | 0.210 | -0.055 | -0.031 | -0.008 |
| Z-7-tricosene | 0.900 | -0.426 | 0.025 | -0.075 | -0.002 | -0.015 | -0.048 |
| Tricosane | 0.019 | 0.011 | -0.713 | -0.023 | 0.660 | -0.232 | -0.027 |
| Nonacosadiene | -0.023 | -0.074 | -0.274 | -0.306 | 0.009 | 0.860 | 0.293 |

**TABLE S17. Perfuming hybrid females with pure species induces differences in their attractiveness.** All the means (number of females mated) and standard deviations (SD) are based on 6 experimental blocks, each with 50 females. All the means (number of females mated) and standard deviations (SD) are based on 6 experimental blocks, each with 50 females. We performed pairwise comparisons using a Tukey test following a One-way ANOVA. The lower triangular matrix shows the t value from a multiple comparisons of means using Tukey contrasts. The upper triangular matrix shows the P-value associated to the comparison. All p-values were adjusted for multiple comparisons.

| ***D. yakuba* males:** $X^2$=40.654, df = 2, P = 1.486 × $10^{-9}$ | | | | |
|---|---|---|---|---|
| | | Pairwise comparisons | | |
| Female type | Mean (SD) | *D. yakuba* | *D. santomea* | NP |
| F1 hybrid perfumed with *D. yakuba* | 0.458 (0.163) | * | 0.009 | 0.117 |
| F1 hybrid perfumed with *D. santomea* | 0.119 (0.043) | 2.787 | * | 0.009 |
| F1 hybrid perfumed with other F1s | 0.319 (0.010) | 1.621 | 2.711 | * |
| ***D. santomea* females:** $X^2$= 44.608, df = 2, P = 2.059 × $10^{-10}$ | | | | |
| | | Pairwise comparisons | | |
| Male genotype | Mean (SD) | *D. santomea* | *D. yakuba* | NP |
| F1 hybrid perfumed | 0.577 (0.088) | * | 0.002 | 0.003 |

| | | | | |
|---|---|---|---|---|
| with *D. santomea* | | | | |
| F1 hybrid perfumed with *D. yakuba* | 0.130 (0.037) | 3.197 | * | 0.005 |
| F1 hybrid perfumed with other F1s | 0.292 (0.073) | 2.938 | 2.783 | * |
| ***D. simulans* females:** $X^2$= 29.695, df = 2, P = 3.562 × $10^{-7}$ | | | | |
| | | Pairwise comparisons | | |
| Male genotype | Mean (SD) | *D. simulans* | *D. mauritiana* | NP |
| F1 hybrid perfumed with *D. simulans* | 0.488 (0.052) | * | 0.002 | 0.008 |
| F1 hybrid perfumed with *D. mauritiana* | 0.145 (0.086) | 3.102 | * | 0.007 |
| F1 hybrid perfumed with other F1s | 0.367 (0.075) | 2.371 | 2.767 | * |
| ***D. mauritiana* females:** $X^2$ =32.307, df = 2, P =9.653 × $10^{-8}$ | | | | |
| | | Pairwise comparisons | | |
| Female type | Mean (SD) | *D. mauritiana* | *D. simulans* | NP |
| F1 hybrid perfumed with *D. mauritiana* | 0.145 (0.085) | * | 0.002 | 0.007 |

| | | | | |
|---|---|---|---|---|
| F1 hybrid perfumed with *D. simulans* | 0.367 (0.075) | 3.1018 | * | 0.006 |
| F1 hybrid perfumed with other F1s | 0.488 (0.053) | 2.767 | 2.371 | * |

**TABLE S18. Perfuming pure-species females with heterospecifics and hybrids induces differences in their attractiveness.** All the means (number of females mated) and standard deviations (SD) are based on 6 experimental blocks, each with 50 females. We performed pairwise comparisons using a Tukey test following a One-way ANOVA. The lower triangular matrix shows the t value from a multiple comparisons of means using Tukey contrasts. The upper triangular matrix shows the P-value associated to the comparison. All P-values were adjusted for multiple comparisons.

| ***D. yakuba* males:** $X^2$ = 10.505, df = 2, P = 0.005 | | | | |
|---|---|---|---|---|
| Female type | Mean (SD) | Pairwise comparisons | | |
| | | *D. yakuba* | *D. santomea* | Hybrid perfumed |
| *D. yakuba* perfumed with *D. yakuba* | 0.411 (0.142) | * | 0.017 | 0.067 |
| *D. yakuba* perfumed with *D. santomea* | 0.231 (0.066) | 2.209 | * | 0.395 |

| | | | | |
|---|---|---|---|---|
| *D. yakuba* perfumed with *D* F1 hybrid | 0.265 (0.074) | 1.920 | 0.851 | * |
| ***D. santomea*** **females:** $X^2$ = 14.813, df = 2, P < 0.001 | | | | |
| Female type | Mean (SD) | Pairwise comparisons | | |
| | | *D. santomea* | *D. yakuba* | Hybrid perfumed |
| *D. santomea* perfumed with *D. santomea* | 0.493 (0.026) | * | 0.002 | 0.003 |
| *D. santomea* perfumed with *D. yakuba* | 0.255 (0.044) | 3.198 | * | 0.965 |
| *D. santomea* perfumed with F1 hybrid | 0.252 (0.048) | 3.183 | 0.096 | * |
| ***D. simulans*** **females:** $X^2$ = 11.409, df = 2, P = 0.003 | | | | |
| Male genotype | Mean (SD) | Pairwise comparisons | | |
| | | *D. simulans* | *D. mauritiana* | Hybrid perfumed |
| *D. simulans* | 0.231 (0.089) | * | 0.005 | 0.002 |
| *D. mauritiana* | 0.306 (0.074) | 2.715 | * | 0.140 |

| | | | | |
|---|---|---|---|---|
| Hybrid perfumed | 0.463 (0.044) | 2.901 | 1.479 | * |
| ***D. mauritiana* females:** $X^2 = 7.3261$, df = 2, P = 0.026 | | | | |
| Male genotype | Mean (SD) | Pairwise comparisons | | |
| | | *D. mauritiana* | *D. simulans* | Hybrid perfumed |
| *D. mauritiana* | 0.463 (0.044) | * | 0.006 | 0.0023 |
| *D. simulans* | 0.306 (0.074) | 2.715 | * | 0.15 |
| Hybrid perfumed | 0.231 (0.089) | 2.9009 | 1.479 | * |

### SUPPLEMENTARY FIGURES

**FIGURE S1. No effect of the marking scheme I. Abdominal color.** Marking females with different abdominal colors had no effect on their attractiveness to pure species males. Confidence intervals around the estimated proportion were calculated using the log (cloglog) parameterization. The red dashed line shows the expectation of completely uniform mating. Nc: non-colored. Table S3 shows the results from the comparisons

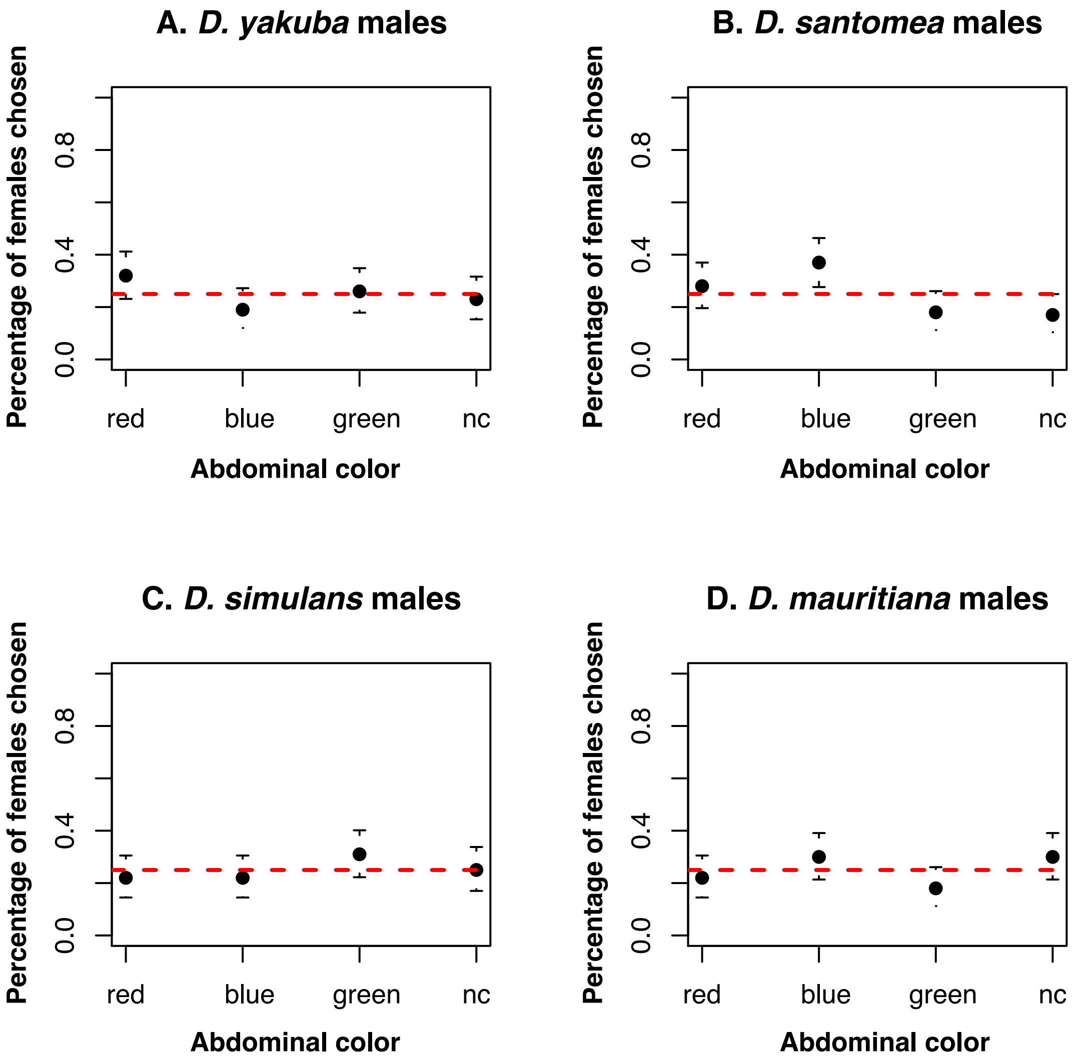

**FIGURE S2. No effect of the marking scheme II. Wing clipping.** Marking females by clipping their wings had no effect on their attractiveness to pure species males. Confidence intervals around the estimated proportion were calculated using the log (cloglog) parameterization. The blue dashed line shows the expectation of completely uniform mating. vert: vertical; hori: horizontal. Table S4 shows the results from the comparisons

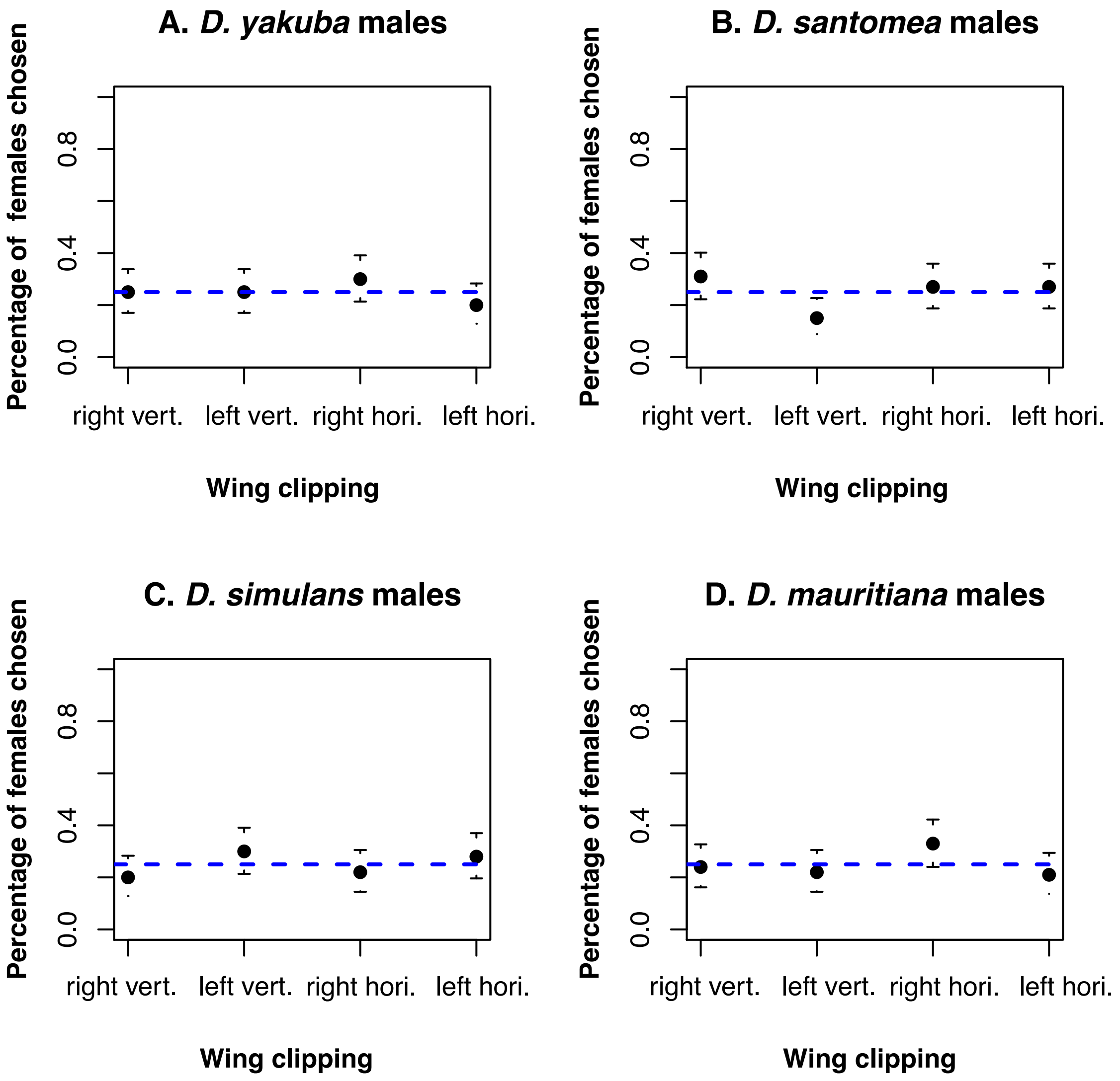

**FIGURE S3. No effect of the marking scheme. Time spent courting each type of female.** blue.l_h: blue, horizontal clipping left wing, green.l_h: green, horizontal clipping left wing, nc.l_h: non-colored, horizontal clipping left wing, red.l_h: red, horizontal clipping left wing, blue.l_v: blue, vertical clipping left wing, green.l_v: green, vertical clipping left wing, nc.l_v: non-colored, vertical clipping left wing, red.l_v: red, vertical clipping left wing, blue.r_h: blue, horizontal clipping right wing, green.r_h: green, horizontal clipping right wing, nc.r_h: non-colored, horizontal clipping right wing, red.r_h: red, horizontal clipping right wing, blue.r_v: blue, vertical clipping right wing, green.r_v: green, vertical clipping right wing, nc.r_v: non-colored, vertical clipping right wing, red.r_v: red, vertical clipping right wing.

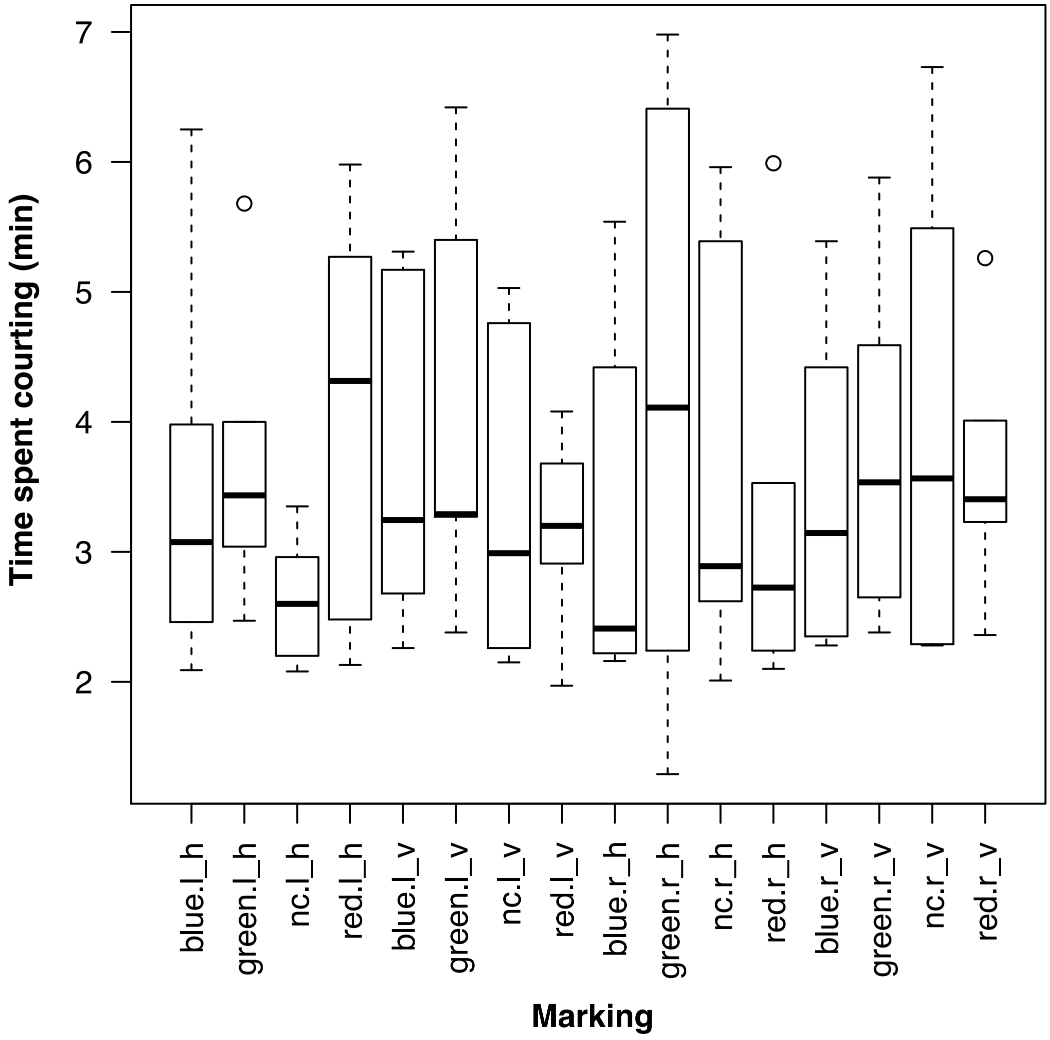

**FIGURE S4. Regression curves for seven CHCs. Each panel shows one CHC.** Linear regressions were fitted through the origin. AUC: Area under the curve.

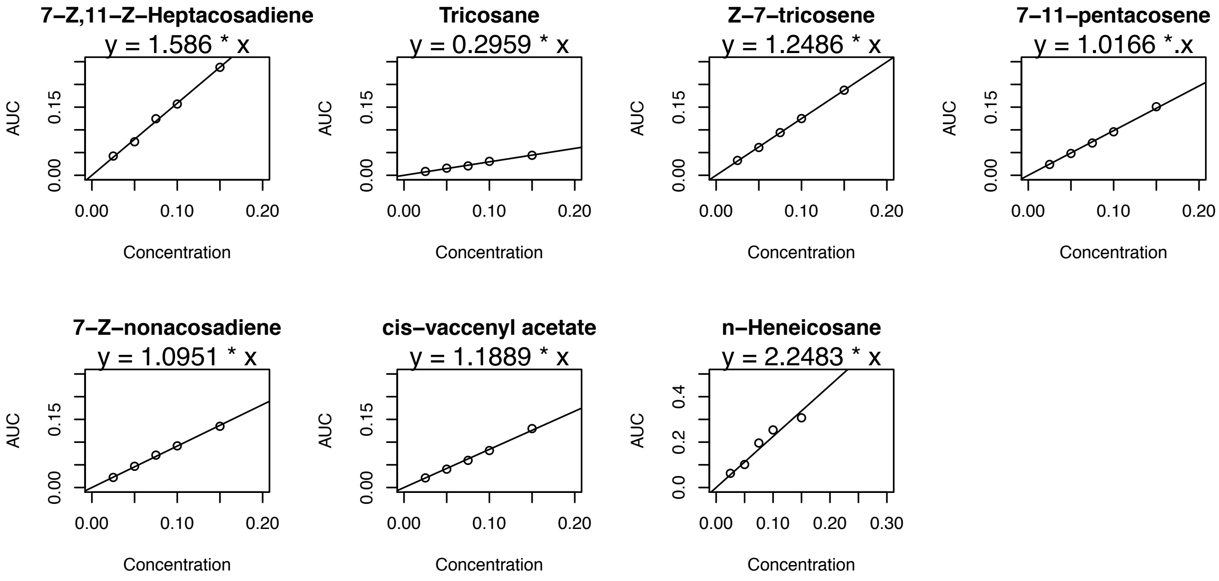

**FIGURE S5. No strong effect of the scorer on the proxy of female attractiveness.**
Each experiment included four observations (i.e., four different females).

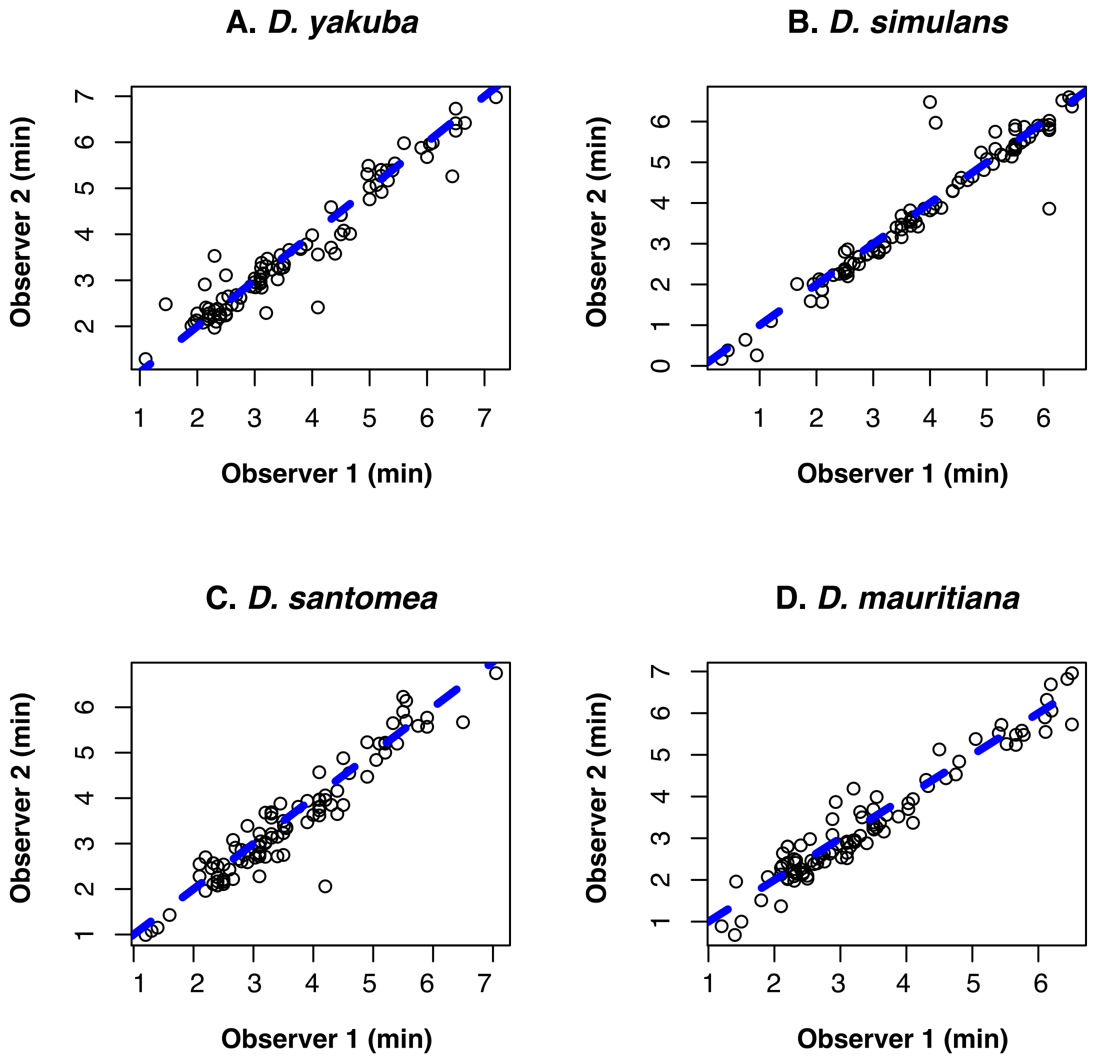

**FIGURE S6. PC1 and PC2 eigenvectors for the *D. yakuba/D. santomea* biplot shown in Figure 4A.**

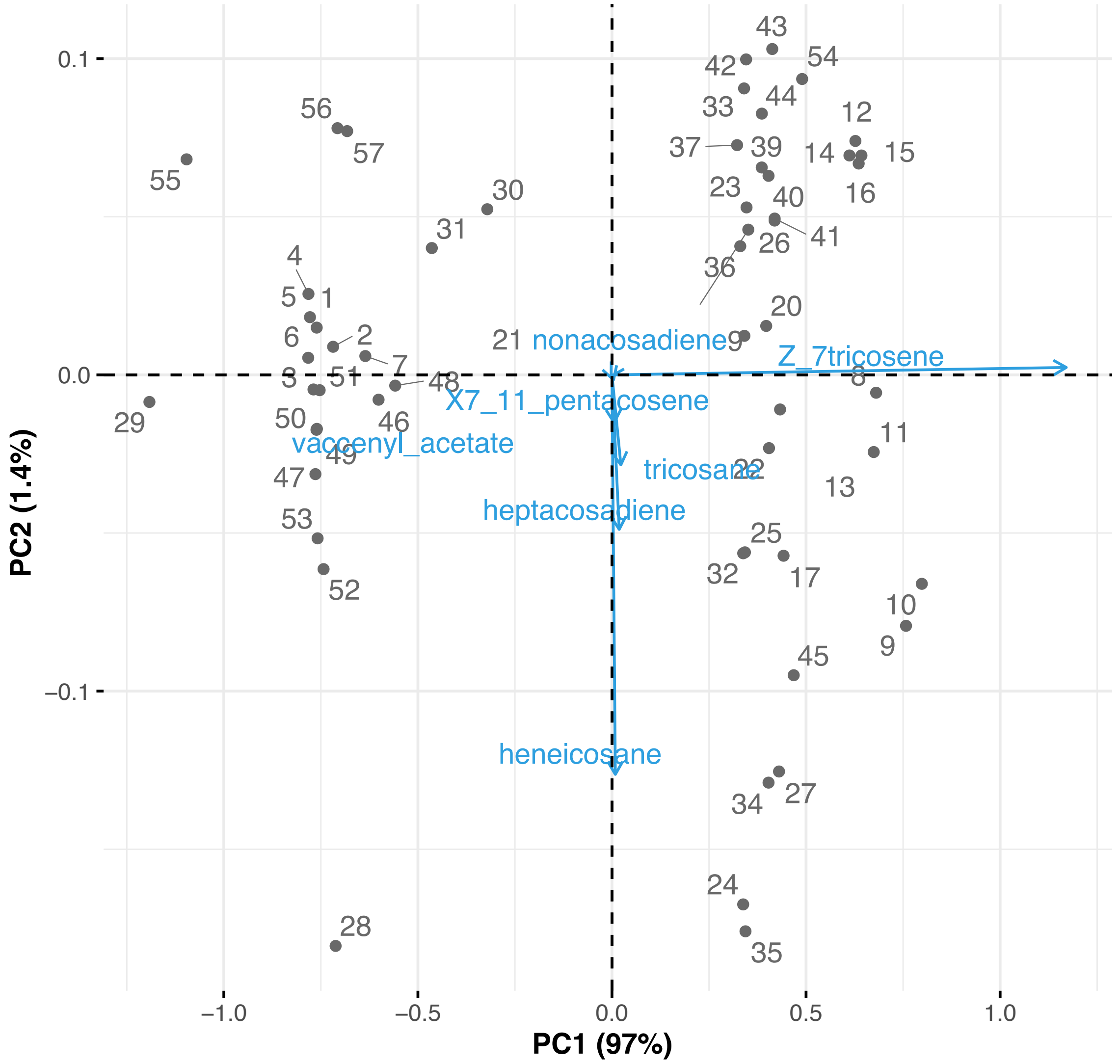

**FIGURE S7. PC1 and PC2 eigenvectors for the *D. simulans/D. mauritiana* biplot shown in Figure 4B.**

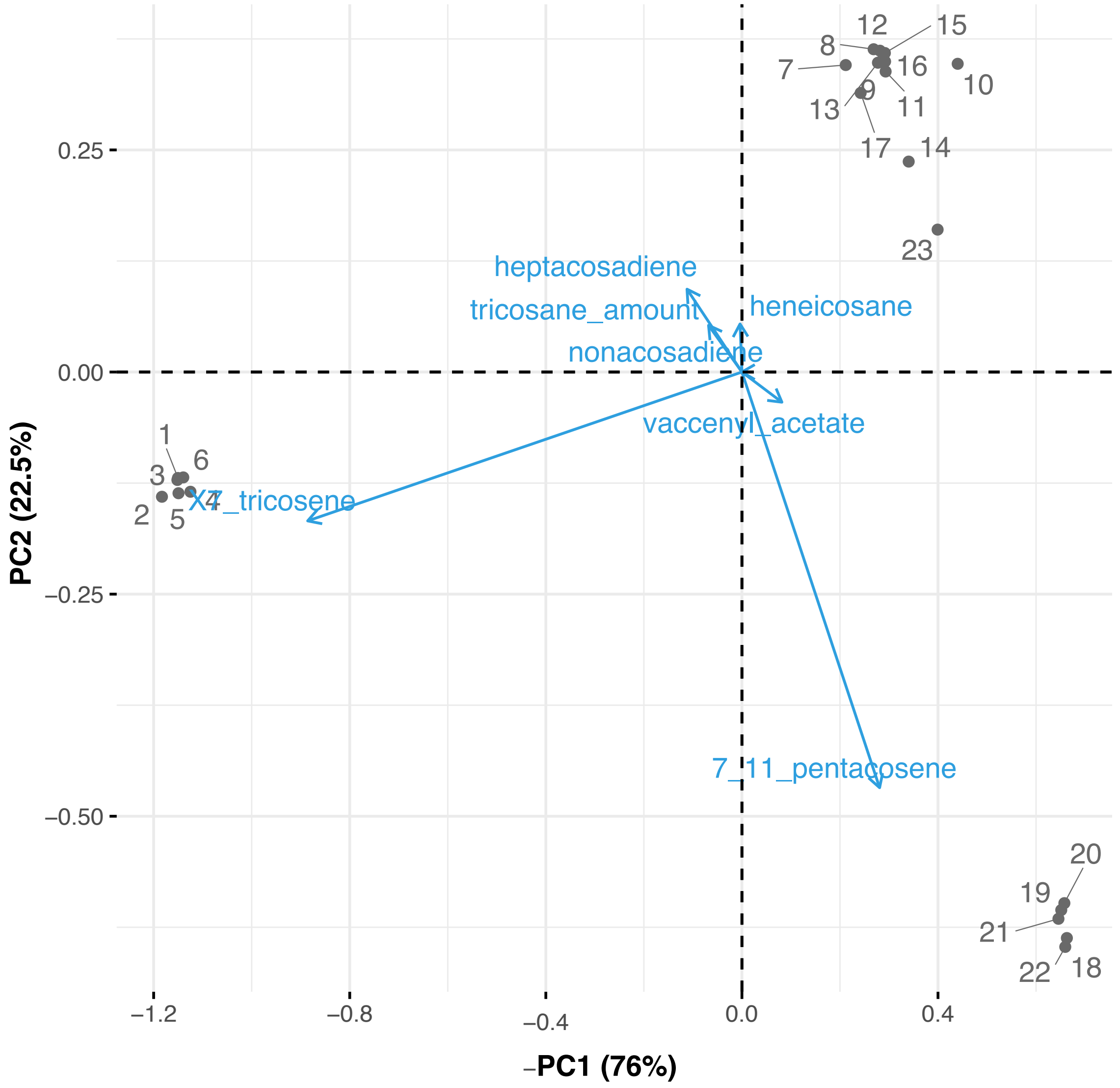

**FIGURE S8. PC1 and PC2 eigenvectors for the perfumed *D. simulans* samples biplot shown in Figure 5A.**

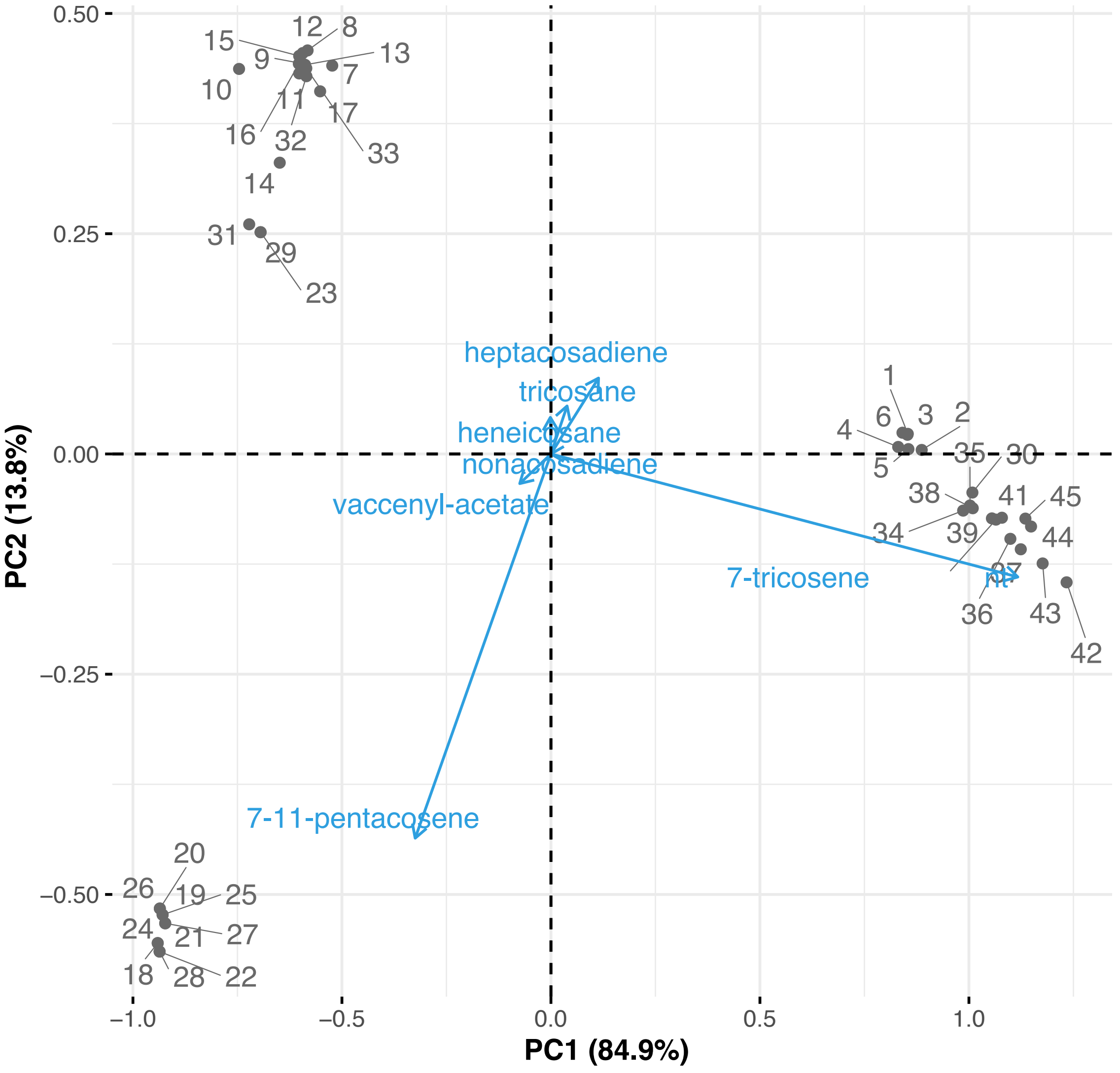

**FIGURE S9. PC1 and PC2 eigenvectors for the perfumed *D. mauritiana* samples biplot shown in Figure 5B.**

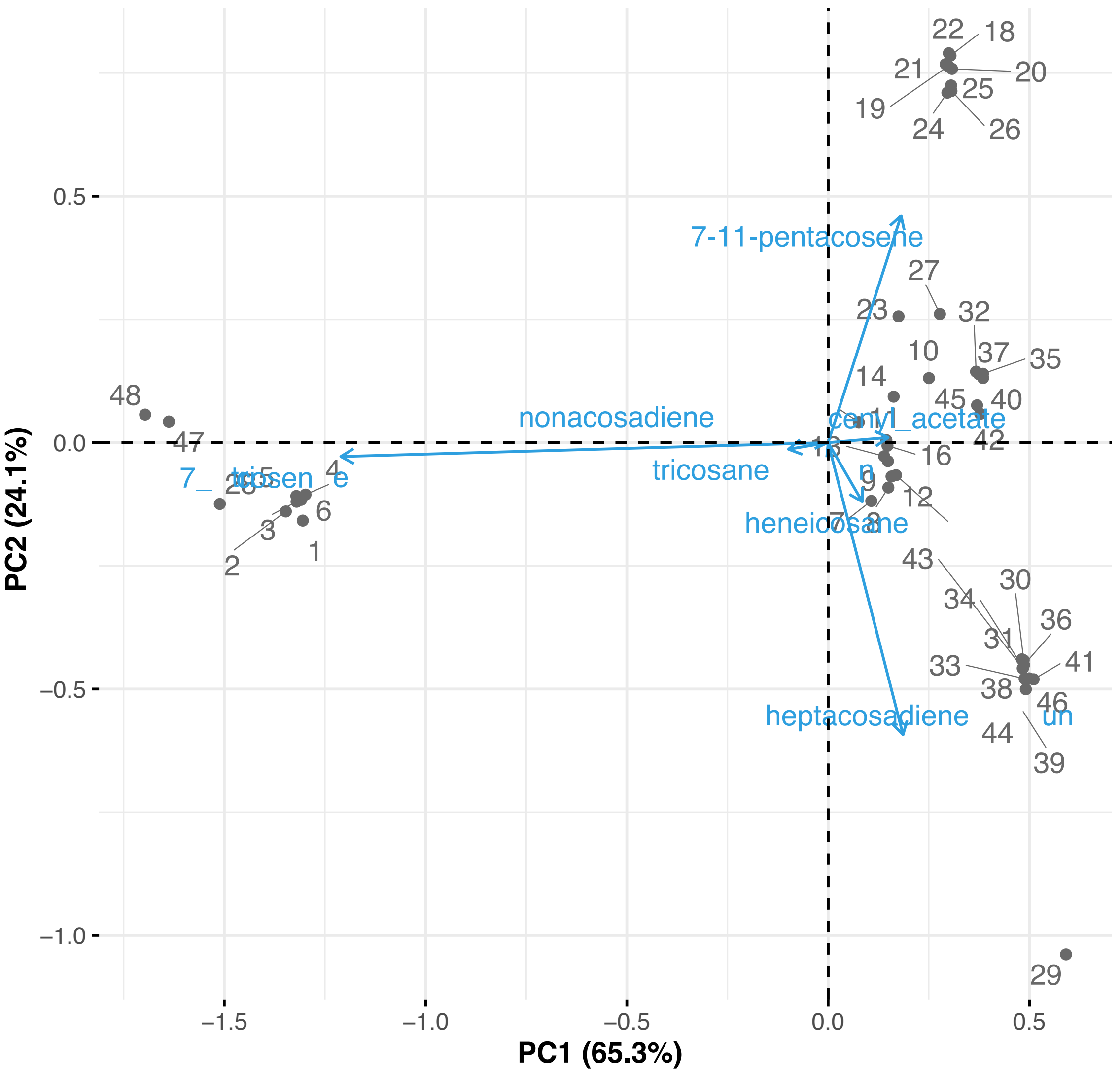

**FIGURE S10. PC1 and PC2 eigenvectors for the perfumed F1 *(sim/mau)* samples biplot shown in Figure 5C.**

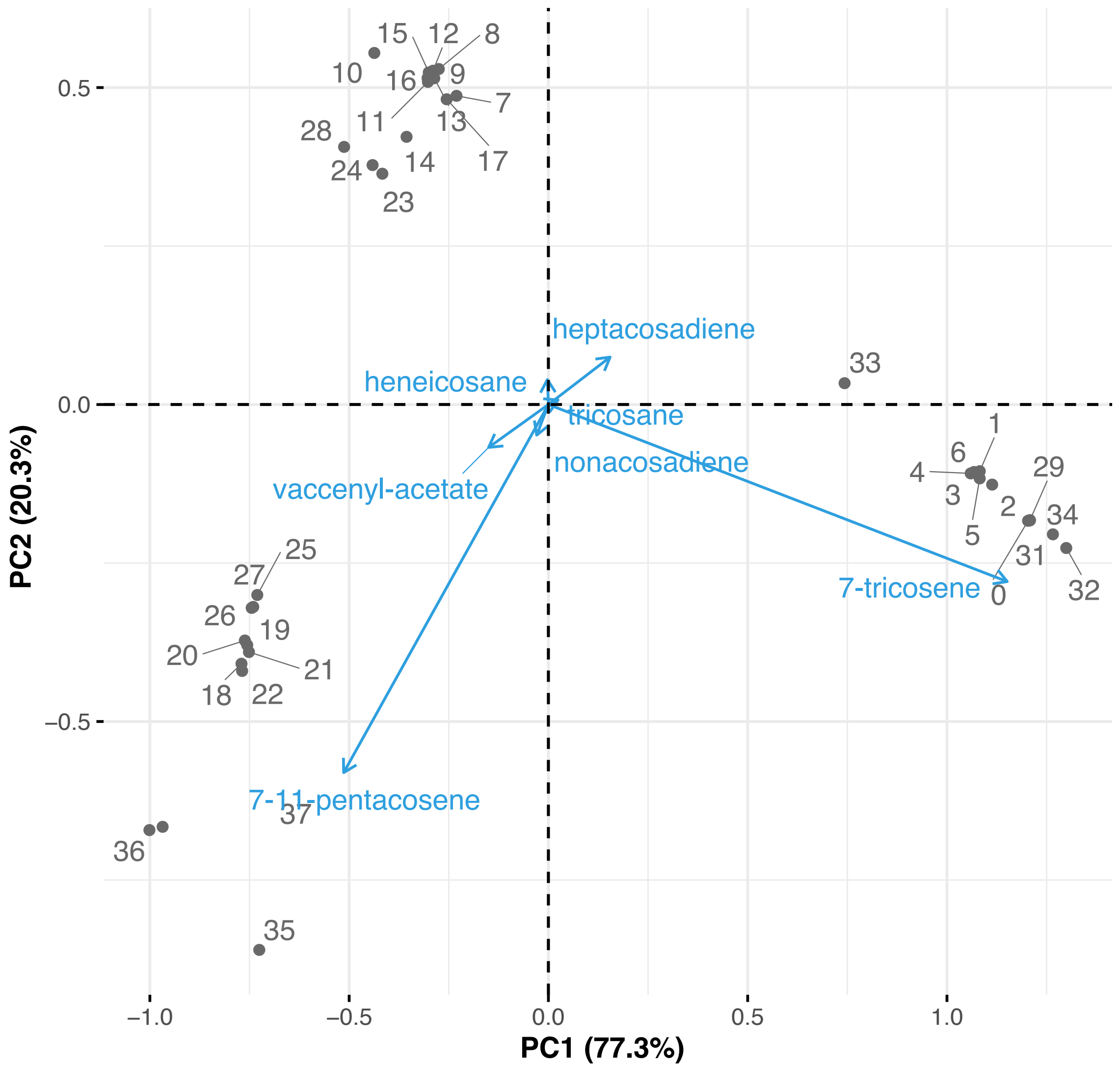
